# Label-Free, Real-Time Monitoring of Cytochrome C Responses to Drugs in Microdissected Tumor Biopsies with a Multi-Well Aptasensor Platform

**DOI:** 10.1101/2024.01.31.578278

**Authors:** Tran N. H. Nguyen, Lisa Horowitz, Timothy Krilov, Ethan Lockhart, Heidi L Kenerson, Raymond S Yeung, Netzahualcóyotl Arroyo-Currás, Albert Folch

## Abstract

Functional assays on intact tumor biopsies can potentially complement and extend genomics-based approaches for precision oncology, drug testing, and organs-on-chips cancer disease models by capturing key determinants of therapeutic response, such as tissue architecture, tumor heterogeneity, and the tumor microenvironment. Currently, most of these assays rely on fluorescent labeling, a semi-quantitative method best suited to be a single-time-point terminal assay or labor-intensive terminal immunostaining analysis. Here, we report integrated aptamer electrochemical sensors for on-chip, real-time monitoring of increases of cytochrome C, a cell death indicator, from intact microdissected tissues with high affinity and specificity. The platform features a multi-well sensor layout and a multiplexed electronic setup. The aptasensors measure increases in cytochrome C in the supernatant of mouse or human microdissected tumors after exposure to various drug treatments. Since the aptamer probe can be easily exchanged to recognize different targets, the platform could be adapted for multiplexed monitoring of various biomarkers, providing critical information on the tumor and its microenvironment. This approach could not only help develop more advanced cancer disease models but also apply to other complex *in vitro* disease models, such as organs-on-chips and organoids.

## INTRODUCTION

As opposed to a modern car or airplane whose state is monitored continuously with hundreds of sensors, most cancer disease models and drug testing platforms do not yield real-time information. In general, information about the state of the tumor is obtained by fluorescent reporters in end-point assays. This shortcoming is especially limiting when the cancer model – typically an animal or a tissue construct – is being exposed to a tumor-killing drug because the drug’s pharmacodynamics can have profound effects on its efficacy and toxicity, as well as its ability to amplify its action with other drugs.^1,2^ While many techniques exist to image tumors *in vivo*, they require complex instruments such as MRI or two-photon microscopes that are not widely available in most laboratories.^3^ Microfluidic electrochemical aptasensors for *in vivo* continuous drug monitoring (MEDIC) have been demonstrated to operate without external reagents or imaging equipment, work at room temperature, and can identify various target molecules by exchanging probes.^4–7^ However, their use as *in vitro* cancer drug probes has only been explored with cell lines and without reporting responses over long periods.^8–10^ Obtaining real-time information about the drug’s efficacy on 3D tumor tissue could help provide more precise information on drug effects and insight into the mechanism of action.

*In vivo* cancer drug testing models result in high attrition rates during clinical development, a failure attributed to inappropriate correlation between the pharmacokinetic and pharmacodynamic parameters, and subsequent extrapolation to human subjects.^11^ In the last decade, *in vitro* models such as patient-derived organoids^12^ and organs-on-chips^13^ have brought some hope for the development of more precise disease models and more efficient drug testing systems, offering simplicity, scalability, and reproducibility. However, in these platforms, which often take weeks to months to establish, the tumor tissue is generated *de novo* from the patient’s cancer cells^14^ or utilizes a synthetic extracellular matrix, generating *in vitro* cultivation bias. These simplified models lack the 3D structural complexity and the molecular and cellular diversity of the human tumor microenvironment (TME),^15^ namely heterogeneous distributions of extracellular matrix (ECM) scaffolds,^16^ immune cells,^17,18^ vasculature, and metabolic gradients^17–19^ that can each impact the tumor’s response or resistance to cancer therapies. Due to these inefficiencies, only a small percentage (∼14%) of the roughly 1,000 drugs entering clinical trials each year manage to meet the criteria for safety and efficacy, a number that drops to less than 4% for cancer drugs.^20^

Researchers have devised functional platforms for assessing drug responses in live tumor samples. Tumor spheroids, also known as "organoids,"^21–31^ formed from patient-derived, dissociated cells, show promise in replicating *in vivo* drug responses due to their 3D architecture that recapitulates some cellular and molecular relationships in the TME.^32^ However, the steps to create and expand tumor spheroids result in the loss of many native immune cells and their original 3D relationships within the TME, limiting their relevance, especially as models for immunotherapy, since many immunotherapies act on the local TME.^33–35^ Microdissected tumors, derived from cutting tumors into submillimeter tissue pieces, maintain the original TME relatively intact.^23,28–31^ Implantable or needle microdelivery devices^36,37^ locally deliver small doses of drugs to the tumor in vivo, with maximal preservation of the TME. Still, issues of tumor accessibility and patient safety limit the applicability of implantable approaches. Patient-derived xenograft (PDX)^38^ mouse models permit the study of drug responses in an intact organism (including immune checkpoint blockade in humanized PDX), but with the caveat that all or most of the TME is from the host mouse and PDX from individual patients grow too slowly to inform initial post-operative therapeutic decisions. Thus, microdissected tumor biopsies with intact TME can potentially complement and extend genomics- and organoid-based approaches for cancer disease models and drug testing by capturing key determinants of therapeutic response, such as tissue architecture, tumor heterogeneity and the TME.^39^ We have recently developed a microfluidic platform based on regularly sized, cuboidal-shaped micro-dissected tumor fragments (referred to as "cuboids") that are mechanically cut with a tissue chopper. More than 10,000 cuboids (∼400 μm-wide) can be produced from ∼1 cm^3^ of solid tumor. The cuboids are never dissociated and retain much of the native TME,^40^ including viable immune cells and vascular structures.

However, despite the great potential of these functional testing platforms for broad disease modeling and drug screening applications, accurately characterizing the behaviors of microdissected tumors and their interactions with drug compounds can be challenging. The levels of secreted biomarkers from tissues serve as key indicators of their dynamic status, changing as they grow, mature, and undergo external stimuli or damage.^41–44^ A significant challenge of current functional assay platforms lies in their limited ability to collect real-time data about the tumor’s secretome as an assessment for drug screening and probing disease pathologies.^45^ At present, most of these assays rely on off-chip or fluorescent labeling analysis, including enzyme-linked immunosorbent assay (ELISA) and Luminex assay (which are quantitative methods best suited to single time-points) or immunostaining analysis (a terminal, labor-intensive, and poorly quantitative assay).^45,46^

Aptasensors employ nucleic acid aptamers to directly measure ligand binding. The aptamers undergo reversible conformational changes when binding to their molecular targets. The electrochemical signal response reflects changes in distance between an electrochemical reporter covalently linked to the aptamer and the gold sensor surface upon binding. This biosensor transforms aptamer-target interactions into an electrically measurable signal, enabling cost-effective, on-chip monitoring of specific biomarkers. Aptasensors offer distinct advantages, as they eliminate the need for processes like washing, separations, chromatographic steps, or costly equipment.^47^ Moreover, this technology has been adapted to monitor drug levels^4,5,48^ or relevant biomarkers, such as cytokines^49^ and other physicochemical parameters from tumor functional testing platforms.^46,50^ Additionally, Kinetic Differential Measurement (KDM) techniques have been developed to address baseline drift (crucial for extended measurements in a cultured environment and accurate calibration), enhance the signal-to-noise ratio, and mitigate variations between aptasensors.^4,51^

The integration of biosensors into functional platforms can offer noninvasive, real-time monitoring of relevant biomarkers *in situ*.^41–44^ We integrated an electrochemical aptamer-based sensor (“aptasensor”) into a cuboid platform to monitor the secretion of cytochrome C (CytC), a cell death indicator, in real time. This mitochondrial redox enzyme is released into the supernatant upon apoptotic cell death.^52^ Because the aptamer can be switched to detect any other secreted compound, the approach can be generalized to any other secreted molecule for which there exists an aptamer. Previous studies have also utilized the CytC aptamer for electrochemical sensors to detect cell death, albeit predominantly as a single-time-point measure and in blood serum samples.^53,54^ Our platform allows on-chip real-time monitoring of the dynamic response of cuboids and their intact TME to pharmaceutical compounds, as drugs can trigger acute, chronic, and/or delayed cellular responses from both tumor cells and non-tumor (e.g., immune) cells. This device could help us understand the temporal dynamics of cellular responses to treatments in intact tissue and thereby shed light on predicting and determining the response of cancer to treatments (Fig. 1A). Complementing the platform’s capability, we designed a multiplexer system for a single-channel potentiostat featuring a printed circuit board (PCB) that interfaces with the electrochemical sensor platform. The multiplexer streamlined the process by reducing costs and simplifying the procedure of reading from multiple electrodes, ultimately enhancing throughput, user-friendliness, and cost-effectiveness, which is particularly valuable given the high expense of amplifiers. With our sensor platform, we detected CytC in the supernatant of mouse or human cuboids that had been exposed to drug treatments. This verification demonstrates our proposed platform’s potential for detecting various relevant biomarkers secreted from intact biopsies. In the future, this platform could help better evaluate targeted therapies and immunotherapy combinations with other drugs, an emerging challenge given the vast number of potential drug combinations and the limited resources for testing.

**Fig. 1.**
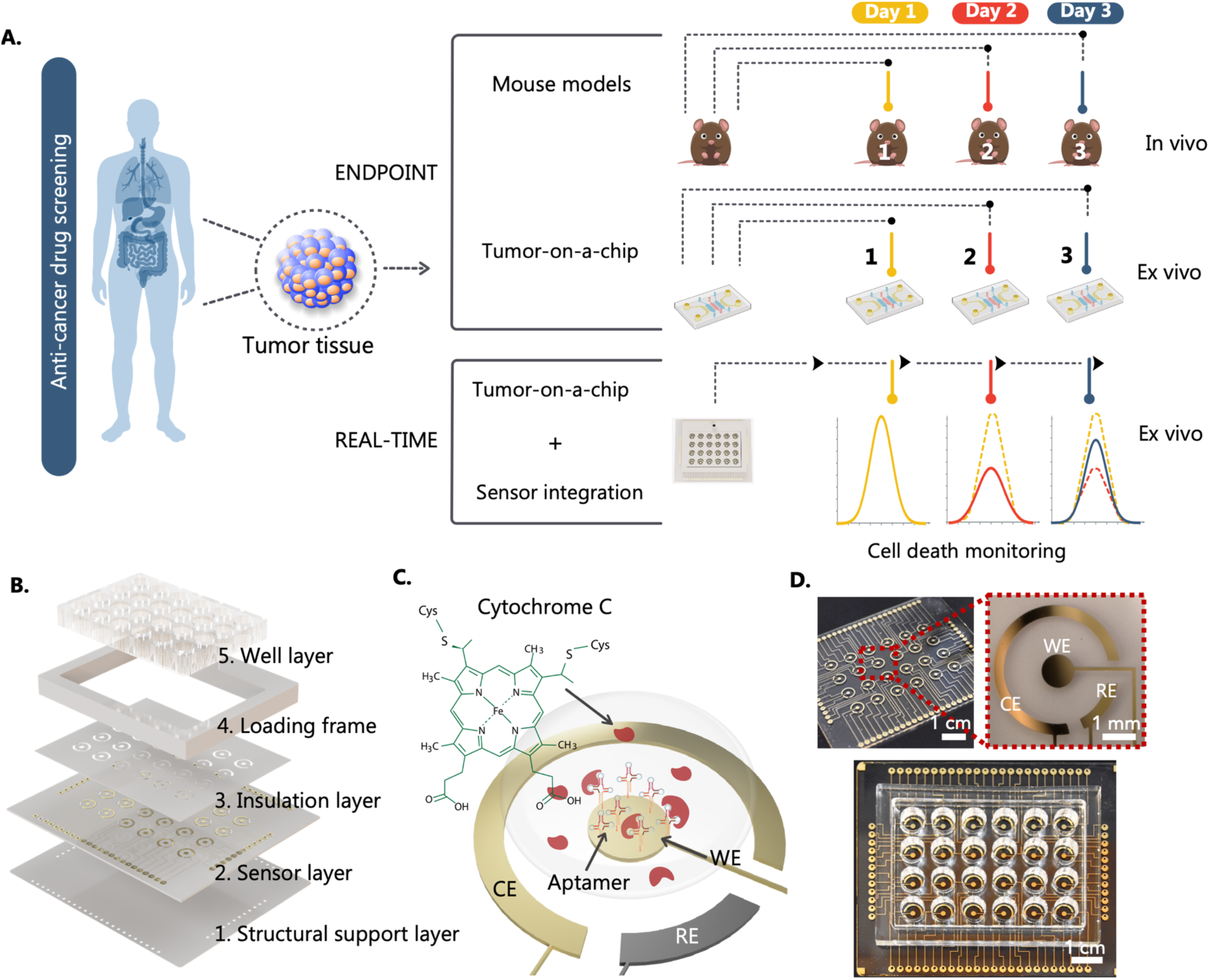
Overview of the electrochemical aptamer-based CytC monitor using microdissected tumor biopsies. **A**. Schematic illustrations of current anti-cancer drug screening models. These approaches predominantly rely on endpoint analysis, providing limited real-time assessments of microdissected tissue responses to drug treatments. Integrating biosensors into the tumor-on-a-chip platforms can enable real-time, on-chip monitoring of the dynamic responses of the tissue of interest to pharmaceutical compounds. **B.** Layer-by-layer design of the sensor platform. Top to bottom: The wells layer, loading frame, and insulation layer define the sensing area, a sensor layer, and a structural support layer. The sensor layer is fabricated using conventional lithography techniques on PMMA. **C.** Schematic illustration of CytC sensing by an electrochemical aptamer-based sensor with the reference electrode (RE), counter electrode (CE), and working electrode (WE) functionalized with CytC aptamers. CytC-induced conformation change of the aptamers causes current response for cell-death detection. **D.** Photograph of the CytC sensor platform. Top left: A photo of the sensor layer on PMMA. Top right: A photo of an individual three-electrode electrochemical sensor with WE, RE, and CE. Bottom: A photo of an assembled CytC sensor platform (Scale bar: 1 cm).

## RESULTS

### Design and fabrication of the multi-well platform with an integrated microelectrode sensor array

We designed and fabricated a custom multielectrode cell culture plate with integrated aptasensors. The custom plate consisted of 24 wells, mirroring the format of a quarter of a standard 96-well plate. It featured 6 columns and 4 rows, with a well-to-well spacing of 9 mm. We used methods adapted from our previous microfluidic multi-well platform for drug testing of cuboids.^55^ The platform (Fig. 1B) is made in poly (methylmethacrylate) (PMMA) layers by digital manufacturing using CO_2_ laser micromachining, solvent bonding, and transfer adhesive techniques.^40,56,57^ The system contains five PMMA layers: (1) a structural support layer; (2) a sensor layer containing the sensing electrodes (fabricated by a combination of photolithography, dry film photoresist technology, and hot roll lamination) on top of an optional microfluidic channel layer; (3) an insulation layer made of removable 3M PolySil silicon adhesive that surrounds the sensor’s working areas and prevents electrical shorting; (4) a frame that forms an outer border; and (5) a bottomless 24-well plate with removable 3M PolySil silicone adhesive (3M300LSE) for the culture stage. The sensor layer consists of an array with 24 sets of bare gold (Au) microelectrodes. Each well has one set of three electrodes: a working electrode (WE), a reference electrode (RE), and a counter electrode (CE). We detail the sensor fabrication process in the Supplementary Materials (Fig. S1). For the base of the aptamer working electrode, we either used bare Au or formed another layer of AuNPs. For the counter electrode, we used a bare Au electrode directly. For the reference electrode, we added another layer of Ag/AgCl. In the last step, we immobilized CytC aptamers onto the working surface using thiol bonds (Fig. 1C-D).

### Characterization of the electrochemical aptamer-based sensor

We developed an electrochemical sensor with a modified aptamer that recognizes CytC, enabling real-time monitoring of CytC increases in the supernatant due to cell death in cuboids during drug screening. We modified the aptamer sequences with a redox-active molecule, methylene blue (MB), at the distal end of the aptamer. As shown in Fig. 2A, the binding of the CytC target induces a rapid and reversible conformational change in the aptamer, increasing the electron transfer (eT) rate between the MB reporter and the WE surface.^5^ This change in eT results in a measurable variation in redox current, easily detectable using square wave voltammetry (SWV). We monitor changes in peak current height, which correlates with the proximity of the MB redox tag to the electrode. The performance of our aptasensors in response to varying concentrations of CytC (ranging from 1 to 30,000 ng/mL) is presented in Fig. 2B. At a recording frequency of 50 Hz, the peak current height of the sensors decreases as the CytC concentration increases. As shown in Fig. 2C, a calibration curve measured at 50 Hz allowed us to estimate an apparent dissociation constant, *K*_D_, of 6.19 ± 1.57 ng/mL (515 ± 130 pM) through nonlinear regression fitting to the Langmuir-Hill isotherm. This *K*_D_ of ∼6 ng/mL, with a dynamic range up to 39,580 ng/mL, indicates the strong binding affinity of our aptasensors for CytC and its ability to detect relevant concentrations (up to 3.32 μM).

**Fig. 2.**
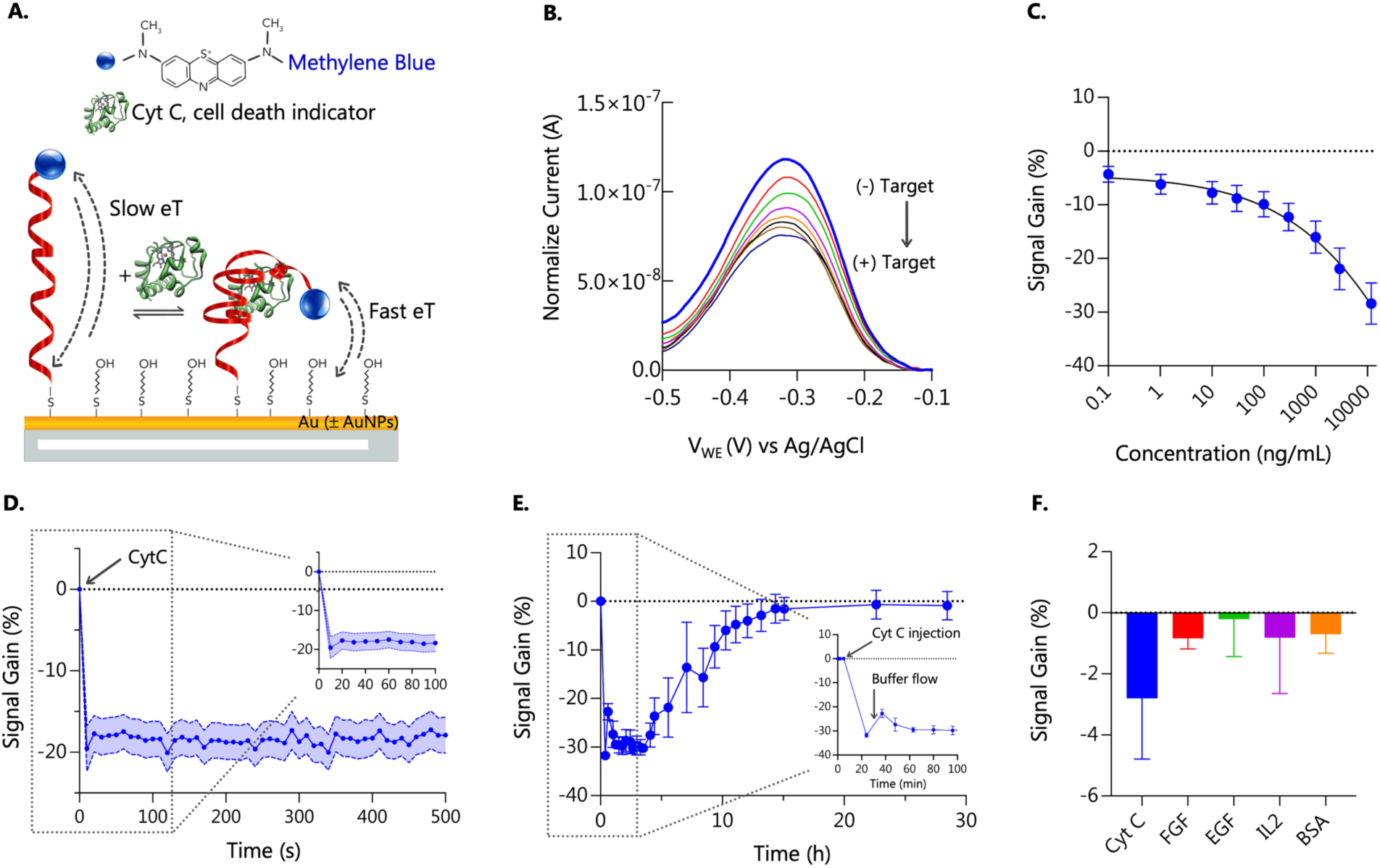
Characterization of the electrochemical aptamer-based CytC sensor. **A.** Aptamer working mechanism. Aptasensors comprise a redox reporter, methylene blue, and hexanethiol-modified aptamer bound to bare Au or AuNPs. In the presence of the target, CytC, aptasensors undergo a binding-induced conformational change, resulting in an electron transfer (eT) difference between the reporter and the electrode. This change can be quantified using square wave voltammetry (SWV). **B.** Square wave voltammogram of CytC aptasensor. These curves represent the currents obtained during SWV scans conducted from -0.5 V to 0 V, utilizing a frequency of 50 Hz and an amplitude of 25 mV. A concentration range of CytC from 0 to 30,000 ng/mL was used (arrows indicate an increase in concentration). **C.** Corresponding dose-response curve of CytC. Error bars indicate the standard deviation (n = 4). **D.** Temporal response to CytC from the aptasensor. Following the addition of 1,000 ng/ml, the CytC aptasensor responds to its target within 100 s (at a SWV of 50 Hz). The shade indicates the standard deviation (n = 3). **E.** Signal responses to CytC injection over time. Initially, the aptamer remained unbound when there was no target (baseline level). Subsequently, as we introduce the target, the association phase begins, and the aptamer binding sites gradually become occupied until they reach a steady-state or equilibrium phase. We then employed continuous buffer flow to facilitate target removal, inducing the dissociation of the aptamer-target complex and returning the signal to baseline levels. Error bars indicate the standard deviation derived from ten scans for each time point. The experiment is duplicated to confirm the results. **F.** Selectivity of CytC aptasensors with response to 100 ng/mL of CytC vs. response in the presence of relevant interferents in medium. Human fibroblast growth factor (FGF), epidermal growth factor (EGF), Interleukin-2 (IL-2), and bovine serum albumin (BSA). Error bars indicate the standard deviation (n = 5).

### Binding kinetics of the CytC aptasensor

We further characterized the binding kinetics of aptamer-protein complexes. Studies looking at the signal in response to a constant amount of CytC ligand indicated that the aptamer-target binding required at least 10 sec to establish equilibrium (Fig. 2C). Next, we determined the monomolecular dissociation rate constant (*k*_off_). Similar to a previous approach to studying *k*_off_ for drug aptasensors,^48^ we adapted our aptasensor platform for flow injection analysis under constant flow. We modified the sensor platform to incorporate an inlet and outlet loop injection system (as shown in Fig. S2A-B). With this setup, we could measure the different phases of aptamer-target interaction, including association, dissociation, and equilibrium. Fig. 2E shows the current response during the injection of the targets. Initially, the aptamer remained unbound in the absence of the target, producing baseline levels of current. The introduction of the target initiated the association phase, which led to the occupation of aptamer binding sites. This phase continued until a steady-state or equilibrium point was attained. Subsequently, the semi-automated target removal initiated the dissociation phase of the aptamer-target complex.^58^ The continuous flow of the medium enabled the aptasensor to recover eventually, gradually returning to the baseline levels within approximately 12 hrs.

### Selectivity of the CytC aptasensor

Selectivity is crucial for the practical application of these sensors in tumor-on-a-chip models for drug screening. To assess the selectivity of our sensors, we conducted experiments in which we measured the sensor’s responses to CytC (100 ng/mL) vs. responses in the presence of other potentially interfering molecules (also at 100 ng/mL). Measurements were performed in a growth medium at 50 Hz. Under these conditions, CytC leads to a higher signal gain compared to others such as human fibroblast growth factors (FGF), epidermal growth factor (EGF), interleukin-2 (IL-2), and bovine serum albumin (BSA) that might be found in culture medium or secreted from tumor tissues (Fig. 2F).

### Kinetic Differential Measurement

To correct for baseline drift (critical for measurement over time in culture and accurate calibration), enhance signal-to-noise, and reduce sensor-to-sensor variability, we applied an approach that uses the difference between signal-ON (increase current) and OFF (decrease current) curves at different frequencies. This approach, called Kinetic Differential Measurement (KDM), was developed previously for *in vivo* electrochemical aptasensors that detect real-time drug levels.^4,51^ KDM measures the signals at two different states of the electron transfer at the WE surface.^59^ At the fast signal-ON state, the electrochemical signal increases with target addition. At the slow signal-OFF state, the electrochemical signal decreases with target addition. Taking the difference between the two measurements, we can improve the gain of CytC sensors and correct the baseline drift (Fig. 3A).^60–62^

**Fig. 3.**
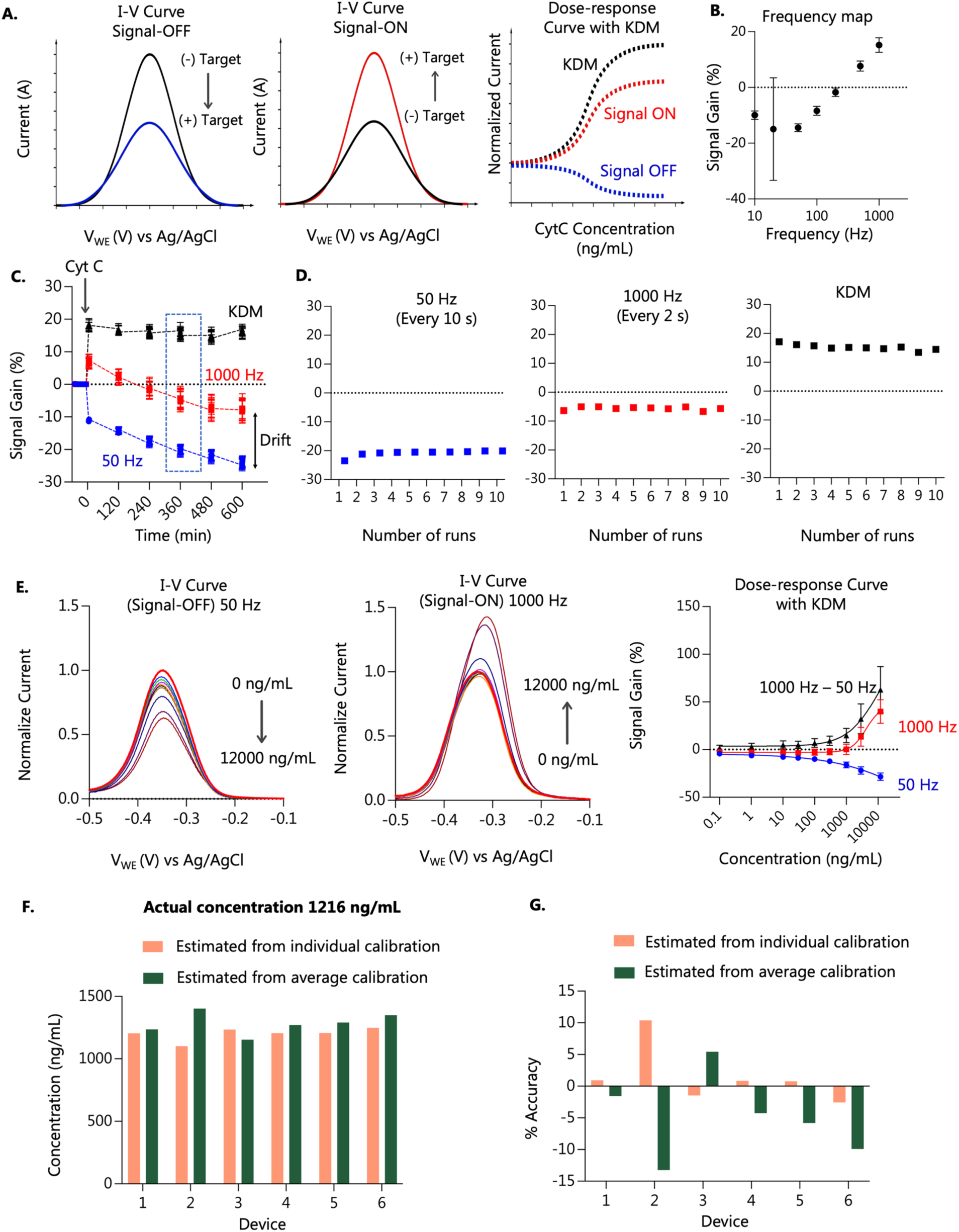
Kinetic Differential Measurement technique and characterization of Au-aptasensor. **A. Model curves illustrating KDM.** The signal gain represents the signal generated when the target concentration increases. Depending on the frequency of square-wave voltammetry used and the sensor’s electron transfer rate, aptasensors can exhibit a decrease in signal gain (Signal-OFF behavior, top left figure) or an increase in signal gain (Signal-ON behavior, bottom left figure) in response to target binding. These characteristics of the aptasensor allow us to calculate Kinetic Differential Measurement (KDM) values to enhance gain and to correct baseline drift. KDM is the difference between the normalized peak currents at Signal-ON and Signal-OFF frequency, divided by the average of Signal-ON and Signal-OFF currents. These KDM values are fitted to the Hill-Langmuir equation to create a dose-response curve (right-side figure). **B.** A frequency map corresponding to different interrogation frequencies measured for CytC-binding by our Au-aptasensors shows dependence on frequency and regions of Signal-On and Signal-OFF frequencies. Error bars represent the standard deviation (n = 5). **C**. **KDM measurement for Cyt C in DMEM-F12-10% FBS culture medium.** KDM (black), as calculated from responses to CytC measured at 50 Hz and 1,000 Hz (blue and red lines, respectively), showed reduced drift over time and increased signal amplitude in a culture medium. Error bars represent the standard deviation (n = 3 sensors). **D.** Expanded graphs of the responses to CytC for one sensor at the 360-minute mark, measured at 50 Hz then 1,000 Hz, and the calculated KDM signal. **E. Responses of CytC aptasensors calibrated in a culture medium.** Response of Au-aptasensors to CytC square wave voltammograms obtained for the 1 – 12,000 ng/mL concentration range at 50 Hz and 1,000 Hz (representative sensor, the arrow indicates an increase in concentration). Graph of KDM values calculated from the normalized peak height at 50 Hz and 1,000 Hz. Error bars represent the standard deviation (n = 12). **F. G. Comparison of actual and estimated concentration from individual sensors using dose-response curve calibration based on KDM values. F.** Estimated concentrations were calculated from individual calibration for each sensor, and estimated concentrations were calculated from average calibration derived from all six sensors. **G.** The percentage accuracy of estimated concentration compared to actual concentration from individual calibration for each sensor and from average calibration derived from all six sensors.

We established the conditions to perform KDM with our CytC aptasensors. Since the relative change of this current is highly dependent on SWV frequency, we interrogated the sensor at multiple SWV frequencies. We probed their amplitude between no target and saturation to generate a frequency map (Fig. 3B). We observed a signal-OFF response from most of the frequencies we tested, from 10 Hz to 200 Hz, and a signal-ON response at 500 Hz and 1,000 Hz. We identified the amplitude-frequency pairs that achieved the most significant signal difference between no target and saturation. As our potentiostat requires more than 50 sec to perform a SWV scan at 10 Hz and generates noisy currents at 20 Hz, we selected 50 Hz for the signal-OFF frequency and 1,000 Hz for the signal-ON frequency for CytC measurements.

Using the previous parameters, we evaluated the stability of KDM recordings. With KDM, we could differentially combine the two output signals to reduce background noise and drift. We exposed the sensors to a CytC concentration of 2000 ng/mL in culture medium (DMEM-F12-10% fetal bovine serum (FBS)) (Fig. 3C) or to Phosphate-Buffered Saline (PBS) (Fig. S3) for 10 hrs. We collected data every 2 hrs with 10 SWV scans each time. The use of KDM reduced the drift when measuring at 50 Hz and 1,000 Hz alone in both cases. Previous studies have indicated two potential distinct mechanisms contributing to signal drift.^51^ The first exponential phase involves fouling by culture medium components. We observed that the drift was more pronounced, as expected, due to fouling from interferents, such as proteins absorbing to the sensor surface, when measuring in the culture medium (Fig. 3C) compared to measuring in PBS (Fig. S3). With KDM, the signal remained stable over 10 hrs in the culture medium. The second phase of signal drift arises from the loss of reporter-modified DNA due to electrochemistry desorption of the monolayer. Thus, there is a tradeoff between the frequency measurements and the duration of the experiment. Therefore, we limited data collection to ten repeated runs every two hours at 50 Hz and 1,000 Hz, for a total of ∼120 runs per sensor. This approach resulted in a stable signal, and any residual drift was effectively corrected using KDM (Fig. 3C-D).

### Optimization of the gold electrochemical aptamer biosensor surface

We evaluated the response of a conventional planar Au surface. The Au-aptasensor setup exhibited a robust signal response, characterized by well-defined MB reduction peaks, across the relevant human CytC recombinant concentration range of 0 – 12,000 ng/mL (Fig. 3E). The Au-aptasensor configuration offered the benefit of straightforward replication and minimized variability, making it the preferable choice for our monitoring objectives. In our exploration of alternative approaches, we considered integrating gold nanoparticles (AuNPs) onto planar Au surfaces before the aptamer immobilization process to enhance the signal-to-noise ratio (SNR) and overall working electrode quality.^63,64^ The AuNPs-aptasensors indeed exhibited higher current responses (Fig. S4A) and displayed a larger-amplitude dose-response curve with average KDM values compared to Au-aptasensors (Fig. S4B-C). However, the fabrication of AuNPs-aptasensors led to significant variability between sensors, rendering it unsuitable for our multielectrode setting. Nevertheless, these AuNPs-aptasensors hold potential for studies requiring fewer sensors and higher sensitivity for measuring lower-abundance biomarkers at low pg/mL levels. Despite the higher sensitivity of the AuNPs-aptasensor, we chose the Au-aptasensor configuration for all experiments due to its suitability for our monitoring requirements, ease of replication, and the assurance of consistent and reliable results, particularly in rapidly detecting CytC at a higher concentration range.

To ensure a high-quality electrode surface, we employed an alternative cleaning protocol, compared to the conventional cleaning method,^4,5^ with hydrogen peroxide (H_2_O_2_) and linear sweep voltammetry (LSV) in potassium hydroxide (KOH), which is compatible with our PMMA-based platform. This method effectively prepared a clean Au surface for subsequent processing, allowing for strong covalent binding of aptamers and efficient signal collection for our aptasensors. The cleaned sensors exhibited distinct MB reduction peaks, indicating the formation of a compact self-assembled monolayer (Fig. S5A). In addition to Au surface cleaning, we addressed the challenge of generating a stable Ag/AgCl film for our on-board reference electrodes. Traditional methods were not feasible due to the PMMA substrate. Instead, we adopted an alternative protocol involving Ag plating, chemical cleansing, cyclic chlorination, improved interfacial adhesion, and a final Nafion layer. This approach provided a thin yet robust Ag/AgCl/Nafion electrode surface, ensuring stability and longevity for our reference electrodes (Fig. S5B). A detailed description of the entire process can be found in the supplementary materials.

### Aptasensor variability

We assessed the variability in sensor performance using KDM. We calibrated six individual sensors at 50 Hz and 1,000 Hz with a concentration range of 0 – 10,000 ng/mL using human CytC recombinant in culture medium (DMEM-F12-10% FBS). Then, we used these calibrated sensors to measure a known concentration of CytC and compared the actual concentrations to the obtained values that were determined from the dose-response curve based on the KDM values. Accuracy, defined as the mean of the relative difference between the estimated and applied concentrations (100 × (expected – observed)/observed) based on a published protocol,^59^ was 1.48% when calibrating each sensor to itself, with a coefficient of variation (100 × population standard deviation / population mean) of 5.28%. However, using an average calibration derived from all six sensors, the mean accuracy slightly increased to 4.29%, but remained within an acceptable range. The coefficient of variation was 6.78%, indicating a marginally increased variability when applying the average calibration (Fig. 3F-G). These results demonstrate low variability within our sensors from the same batch, confirming their suitability for reliable monitoring.

### Automated switching between electrodes using an electronic multiplexer

Electrochemical potentiostats are expensive. To reduce the cost and simplify the procedure of reading from multiple electrodes, we implemented an electronic multiplexer setup that allowed for the automated switching of a single potentiostat to each of the 24 sensors. The reduction in bandwidth does not affect our final measurements because CytC secretion is a slow process. The system included a customized PCB board to plug in the sensors and three 32-to-1 channel multiplexer chips (one for the three sets of WE, RE, and CE electrodes), all situated on a separate PCB board. A flat flex cable (FFC) connected the PCBs and allowed us to separate the sensor platform board as a stand-alone unit for incubation in a 37 °C incubator. Furthermore, we established communication between the electrode array and a single-channel commercial potentiostat (DY-2219, Digi-Ivy) by connecting the multiplexing board with a HiLetgo microcontroller (Fig. 4A). This setup enabled electronic switching between various electrodes and enhanced throughput for drug applications on microdissected tissues, offering significant advantages over single-working electrode-based electrochemical detection platforms. Fig. 4B illustrates the hardware block diagram for the multiplexing platform from the sensor arrays through the multiplexers to the potentiostat. We evaluated the effect of the multiplexer on the electrochemical signal acquired from the sensor platform. We compared SWV results obtained directly through a single-channel potentiostat and the SWV results obtained indirectly through the multiplexer interface. Fig. S6A demonstrates that both conditions exhibited similar responses, with less than 5% difference in normalized peak values at 50 Hz and 1,000 Hz across the tested devices (Fig. S6B).

**Fig. 4.**
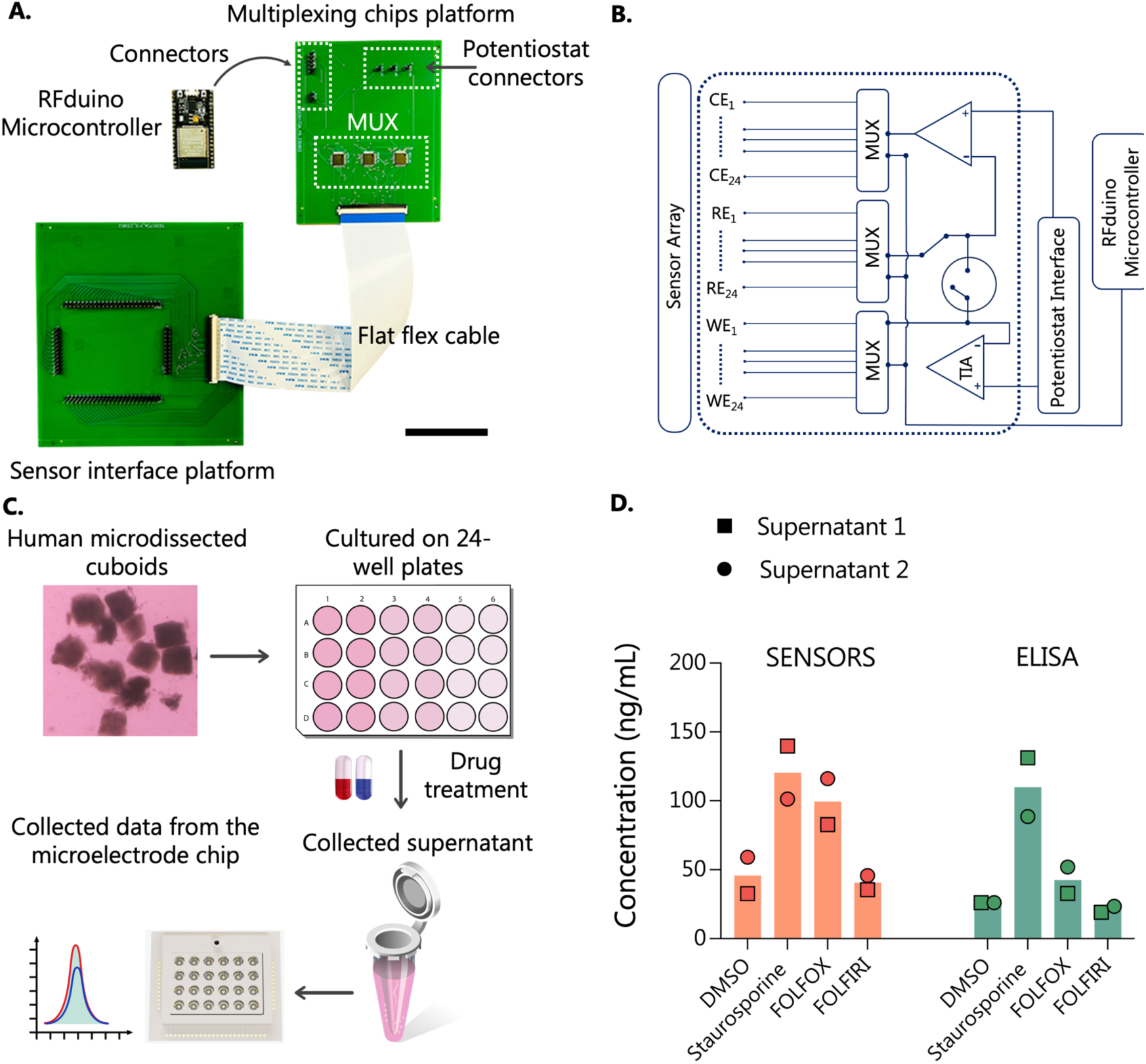
Detection of CytC in the supernatant of treated human cancer cuboids using a multiplexing system. **A.** Photograph of the multiplexing system with PCBs for sensor interfacing and multiplexing chip housing. **B.** Hardware block diagram for the multiplexing platform from the sensors through the multiplexers to the potentiostat. WE1: working electrode 1, RE1: reference electrode 1, CE1: counter electrode 1; MUX: multiplexer. Scale bar, 5 cm. **C.** Schematic illustration of the supernatant CytC sensing experiment, where we cultured cuboids collected from a patient’s tumor in a 24-well plate for 3 days under different drug treatment conditions, with each condition duplicated. We collected and measured CytC from the supernatants using ELISA and an electrochemical aptamer-based sensor platform. **D.** ELISA and electrochemical sensors measurements of CytC from human μDT tissues from two supernatants after treatment for three days with Staurosporine, FOLFOX, and FOLFIRI compared to DMSO control vehicle. Individual data points represent each sensor, with red and blue lines corresponding to duplicate supernatant samples.

### Measurement of CytC from the supernatant of human microdissected cancer tissue cultured with drugs using the aptasensor platform

We conducted a study to assess our aptasensor platform’s ability to quantify CytC secretion from human cuboids after drug treatments for three days in culture with Staurosporine, FOLFOX (a combination of fluorouracil and oxaliplatin), and FOLFIRI (a combination of fluorouracil and irinotecan), in comparison to the dimethyl sulfoxide (DMSO) vehicle control. We prepared uniform-sized cuboids from a human colorectal tumor biopsy using a previously documented procedure.^40^ We performed each drug condition in duplicate. Subsequently, we collected the supernatants and measured the level of CytC released from the cuboids after cell death using our aptasensor platform. To validate our platform’s measurements, we compared the results with those obtained from ELISA (Fig. 4D).

Our aptasensor platform showed a marked increase in the CytC production with Staurosporine treatment (∼3-fold) compared to the DMSO control vehicle. In contrast, compared to the DMSO control vehicle, we observed a more modest increase (about 2.25-fold) with FOLFOX treatment and a slight decrease (0.92-fold) in CytC production with FOLFIRI treatment. The trend from ELISA results closely follows this observation, with CytC production increasing by 4.23-fold for Staurosporine, 1.6-fold for FOLFOX, and a decrease to 0.73-fold for FOLFIRI, all relative to the DMSO control vehicle, based on duplicated supernatant samples. In some cases, our platform even indicated higher production rates. Duplicates for both the sensor and for the ELISA were consistent. Our results aligned overall with those from ELISA, confirming the aptasensor’s capability to measure CytC production from human cuboid samples in real time. However, we observed a slight discrepancy in CytC concentration with FOLFOX treatment. While the trend in measurements for supernatant 1 and supernatant 2 was similar between sensor reading and ELISA, the higher concentration observed in sensor readings may stem from variations in our sensor-specific calibration curves.

### *Ex-vivo* monitoring of CytC from microdissected tumor tissues after drug treatment in culture using the aptasensor platform

We validated our sensor platform’s ability to quantify CytC secretion from cuboids at different time points during an extended culture period (48 hrs). We prepared cuboids from human U87 glioma cell-derived tumors grown in athymic nude mice. After overnight culture in a 96-well plate to allow recovery from tissue damage due to the cutting procedure, we transferred the cuboids onto our sensor platform for measurement.

To prevent random adhesion of the cuboids to the sensor surface and improve the precise localization of CytC secretion detection, we encapsulated the cuboids in a hydrogel. We mixed them with hydrogel before transferring them onto the sensors and immobilizing them to a restricted area just above the working electrode (Fig. 5A-B). The hydrogel matrices exhibited interconnected pores and good water retention, thus maintaining a suitable environment for tissue proliferation and growth. Encapsulation of cells in hydrogels in close proximity to the WE has been shown to generate a more localized and distinguishable electrochemical response.^65^ Hydrogels have been used as a protective layer to ensure the long-term stability of aptasensors.^66,67^

**Fig. 5.**
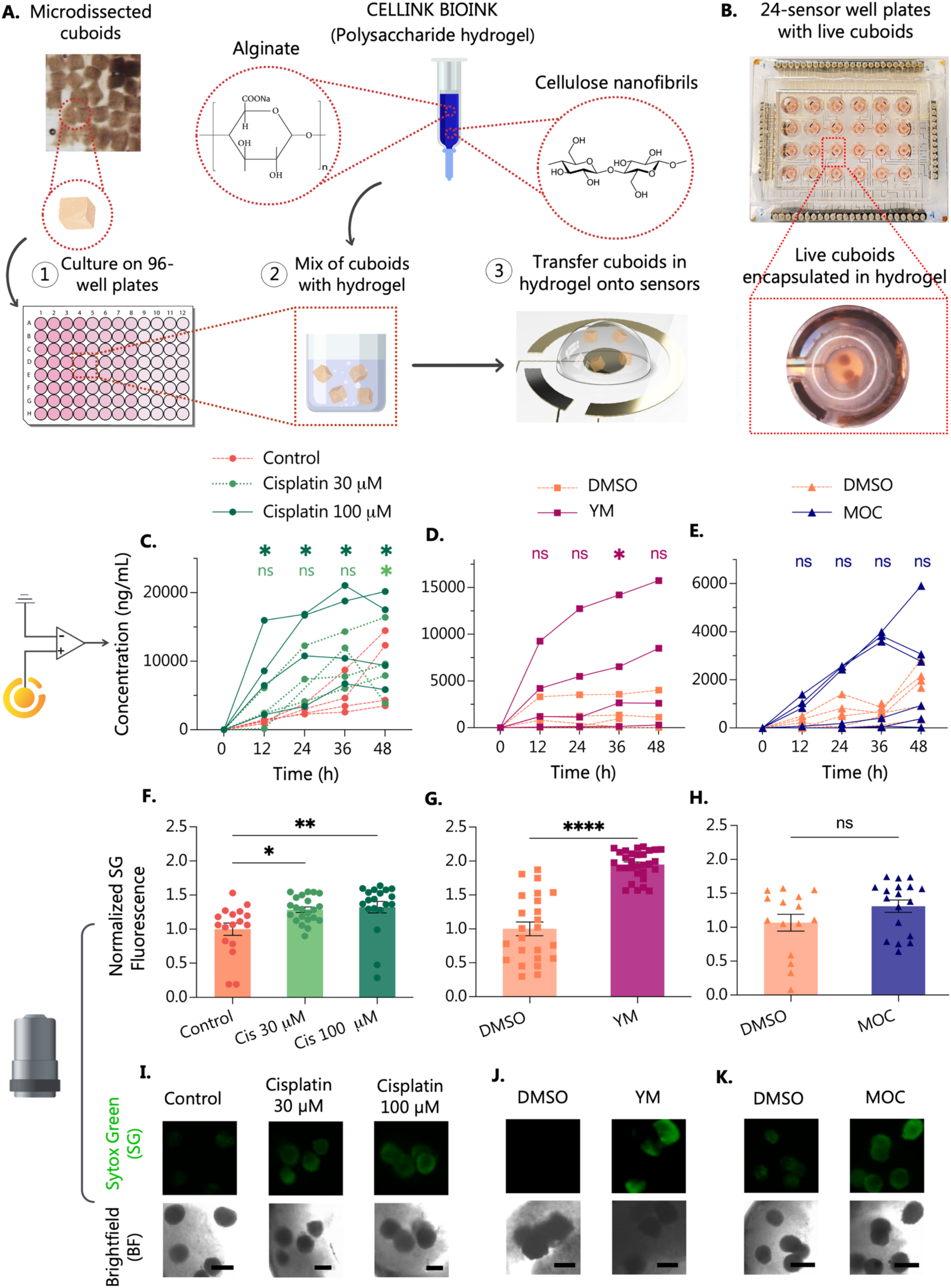
Direct, on-device measurement of CytC secretion from microdissected tumor "cuboids" cultured and exposed to drugs on the electrochemical aptamer-based sensor platform over two days. **A.** Schematic illustration and **B.** photographs of an *ex vivo* experiment, where microdissected U87 glioma xenograft cuboids in Cellink bioink were transferred on the aptasensor system for CytC detection. **C.** Measured CytC concentrations from U87 cuboids after treatment for 48 hrs with different concentrations of cisplatin compared to control. Individual points for every sensor. n = 4. **D.** CytC corresponding concentrations in U87 cuboids after treatment for 48 hrs with YM-155 compared to DMSO control vehicle. Individual points for every sensor. n = 3-5. **E.** CytC corresponding concentrations in U87 cuboids after treatment for 48 hrs with MOC compared to DMSO control vehicle. Individual points for every sensor. n = 5-6. (D-E. Paired t test *versus* control. ns p > 0.05, *p < 0.05). **F-H.** Quantitation of cell death by SYTOX Green (death) fluorescence. Mean fluorescence was normalized to the average value of control conditions. Individual points and average ± S.E.M. n = 25-30. (One-way ANOVA versus control, Dunnett’s multiple comparison test. ns p > 0.05, *p < 0.05, ** < 0.01, **** p < 0.001). **I.** Cell death in U87 cuboids after treatment for 48 hrs with different concentrations of cisplatin compared to control. **J.** Cell death in U87 cuboids after treatment for 48 hrs with YM-155 compared to DMSO control vehicle. **K.** Cell death in U87 cuboids after treatment for 48 hrs with MOC compared to DMSO control vehicle. SYTOX Green (green nuclear death stain), BF (bright field). Representative images from one sensor well with four to five cuboids in each well (Scale bar: 400 μm).

We used a polysaccharide hydrogel (Cellink, Bioink, Boston, USA) that contains nanofibrillar cellulose and sodium alginate. We found that placing this polysaccharide hydrogel atop the working electrode (WE) did not hinder the sensor’s ability to detect CytC and perform KDM (Fig. S7A). We also observed that the performance of the sensors remained unhindered regardless of the hydrogel concentration, apart from an extension of the reaction time to approximately 30 min (Fig. S7B). However, we also found that a 100% hydrogel concentration posed challenges during the transfer of encapsulated cuboids to the surface, potentially resulting in the inadvertent removal of electroactive aptamers during the transfer. As a result, we diluted the hydrogel concentration to 50% with Dulbecco’s Phosphate-Buffered Saline (DPBS), permitting effective transfer while ensuring proper hydrogel adhesion to the electrode surface. We did not use the natural hydrogel, collagen, because its application on top of the electrodes impaired the detection of CytC and did not allow for KDM (Fig. S7C-D).

Next, we evaluated our sensor platform’s ability to measure CytC secretion from cuboids in response to exposure to cytotoxic drugs. Initially, we measured the baseline signal with the encapsulated cuboids on the WE before exposing U87 cuboids to different concentrations of cisplatin (a cytotoxic chemotherapy drug, Fig. 5C), sepantronium bromide, (YM-155, a survivin-inhibitor and potential anti-cancer drug, Fig. 5D), and mocetinostat (MOC, a histone deacetylases inhibitor cancer drug, Fig. 5E) for 48 hrs. For cisplatin, we treated six wells, each at a different concentration (0, 30, 100 μM), and collected the sensor signal every 12 hrs. For YM-155 and MOC, we treated eight wells, each at 1 μM and 10 μM, respectively, compared with DMSO vehicle control and collected the sensor signal every 12 hrs. Untreated cuboids exhibited lower CytC values from 12 to 48 hrs. Tissues cultured with 100 μM cisplatin exhibited the highest levels of CytC, beginning at 12 hrs and showing incremental growth over the 48-hr period. The cuboids cultured with 30 μM cisplatin displayed a higher CytC level compared to the control but with a different time course. CytC levels increased in more pronounced increments from 12 to 24 hrs and 24 to 36 hrs, followed by marginal growth from 36 to 48 hrs. Notably, the result closely resembled the range observed in tissue treated with 100 μM cisplatin at the 48-hr mark.

Additionally, we conducted experiments in which we measured the sensors’ responses to CytC (100 ng/mL = 8.3 nM and 2000 ng/mL = 166 nM) versus responses in the presence of all the drugs used to treat the microdissected tumor tissues, ensuring the selectivity of the sensors in these conditions. We performed measurements in a culture medium at 50 Hz and 1,000 Hz and generated KDM signals from measurements at 50 Hz and 1,000 Hz (Fig. S9A-C). Under these conditions, CytC responses led to a higher signal gain compared to the responses in the presence of relevant drug concentrations, such as DMSO control vehicle (0.2%), 10 μM MOC, FOLFOX (1 μg/mL 5FU + 1 μg/mL Oxaliplatin), 30 μM and 100 μM Cisplatin, 1 μM Staurosporine, and 1 μM YM-155.

We also performed terminal live/dead staining of the cuboids (Fig. 5F-K) to confirm our electrochemical obtained with the sensor platform (Fig. 5C-E). After 48 hrs in culture, we stained the cuboids with the green fluorescent dead nuclear stain, SYTOX green (SG), and the blue fluorescent pan-nuclear stain, Hoechst (H) (Fig. 5F-K). Control cuboids displayed lower SG dead stain fluorescence (normalized to the mean), whereas treated cuboids with 30 and 100 μM cisplatin exhibited higher SG dead stain fluorescence. Our findings indicated a statistically significant response to 30 μM and 100 μM cisplatin (Fig. 5F, I). These observations were consistent with our previous fluorescence-based findings^40^ and agreed with our sensor-based responses. This endpoint live/dead analysis confirmed the CytC aptasensor cell death responses to various cisplatin concentration treatments.

We also performed similar experiments with the CytC aptasensors cuboids treated with other chemotherapy drugs, YM-155 (Fig. 5D) or MOC (Fig. 5E) versus DMSO vehicle control. Similar to cisplatin, we observed that the YM-155 treatment also showed cell death responses by increases in CytC. The treatment with YM 155 exhibited a higher CytC production rate than the DMSO control, first seen at 12 hrs and displaying incremental growth over the 48-hr period. Our terminal live/dead staining revealed elevated SG dead stain fluorescence in cuboids treated with YM-155 compared to the DMSO control (Fig. 5G, J).

Finally, we noted a similar trend with the treatment involving MOC, though at much lower levels and not statistically significant. The MOC-treated cuboids exhibited a higher average CytC production rate than the DMSO control. However, the sensor data indicated that the variance in CytC secretion levels between MOC-treated cuboids and the DMSO control was less pronounced compared to the difference observed in the YM-155 treatment. Our findings were substantiated by our cell live/death staining, which revealed slightly elevated SG dead stain fluorescence in cuboids treated with MOC compared to the DMSO control (Fig. 5H, K).

## DISCUSSION

We have introduced an integrated electrochemical sensor platform that enables on-chip, real-time monitoring of cell death by increases of CytC in the medium of long-term cultures of intact tumor biopsies. We made these measurements with a multi-well sensor layout and a multiplexed electronic setup. With this platform, we showed the *in-situ* response of cuboids to various chemotherapy treatments. The sensors demonstrated selectivity, specificity, and reproducibility while minimizing interference. We have also shown the capability of our aptasensors to effectively utilize Kinetic Differential Measurement (KDM), a technique specific to the aptamer used, to address signal drift caused by long-term culture.

The temporal sensitivity of our CytC aptasensor results from the binding kinetics of the CytC aptamer receptor, with a fast ON response (seconds) and a slow OFF response (hours). A fundamental tradeoff between thermodynamic characteristics (sensitivity) and kinetic properties (time resolution) actively influences molecular interactions.^68,69^ In situations such as ours, with high-affinity receptors and low target concentrations, achieving equilibrium between the receptor’s bound and unbound states takes longer due to a slow *k*_off_ rate.^70,71^ These kinetics introduce a slight delay in generating the final sensor readout. This characteristic is evident in our case, where a higher receptor affinity results in a slower off-rate (*k*_off_ = 12 hrs). This slow timescale of OFF measurements is compatible with our goals for a CytC cell death sensor. Cell death markers like CytC are not continuously released, but released during specific events, such as apoptosis induced by chemotherapy drugs. This transient secretion pattern means that CytC concentration in the culture medium can fluctuate over time. Because of the slow decrease in signal over hours, we can be confident to not miss increases in CytC that may have occurred between readings, which we take periodically. Thus, stable binding to the sensor surface is crucial, allowing us to track dynamic increases in CytC levels while minimizing background noise. We can allow the sensor more time to reach equilibrium to address the slow equilibration kinetics.^68,69^ Consequently, our slow, high-affinity sensor allows us to measure low-abundance analytes in the nM range over hours, compared to other, real-time, small-molecule sensors (e.g. for drugs) that measure higher abundance analytes in the μM range over seconds/minutes.^64,72,73^

Our integrated sensor platform, as demonstrated by *ex vivo* characterization results, effectively enables us to track CytC release over time under drug treatments. The platform reliably yields high SNR ratio measurements during extended *ex vivo* operations (up to 48 hours). Our experiments demonstrated the capability of our Au planar sensors to achieve high SNR measurements of CytC over prolonged periods in the culture medium. Furthermore, the successful fabrication of AuNPs with an increased surface area for aptamer immobilization positions our platform for future applications, particularly in measuring lower-abundance biomarkers like cytokine biomarkers at the pg/mL level.

In this study, we demonstrated the utilization of hydrogel to prevent fouling and improve the precise localization of CytC secretion detection. The choice of hydrogel can affect the performance of the aptasensor. The electrochemical aptasensor relies on biorecognition elements, including the incorporation of a redox probe at the 3’ distal end of the oligonucleotide, and the resulting signal depends on a conformation change of the surface-bound aptamer.^66,74^ As sensor signaling relies on aptamer conformation and flexibility, the aptamers must be able to move freely upon target binding to generate electron transfer between the redox probe and the electrode surface.^60,66^ Hence, it is essential to ascertain that the hydrogel does not appreciably impact the signaling capacity of the sensors and that the sensors can quantitatively respond to CytC at varying frequencies to ensure effective utilization of KDM. We first considered collagen matrices since they are commonly utilized for cell culture and microencapsulation of proteins or small molecules for drug delivery.^65,75^ Unfortunately, collagen encapsulation obstructed the aptasensor’s ability to perform KDM, even at half strength (Fig. S7C). Moreover, collagen led to a prolonged reaction time, preventing the sensor from reaching an equilibrium state even after 90 minutes (Fig. S7D). We hypothesize that the density of the collagen gel partially impairs the conformational change of the aptasensor upon binding. These limitations prompted us to explore alternative hydrogels. One alternative, Cellink bioink, exhibited successful sensor performance (Fig. S7A-B). Although a biophysical characterization of the hydrogels is beyond the scope of the present study, this finding underscores the importance of material selection for sensor signaling capacity.

Despite successful periodic monitoring and discrimination of CytC secretion induced by different drugs and concentrations during extended culture periods, challenges arise in conducting measurements on a large scale and managing variability in drug and culture conditions. We observed a slightly large standard deviation in the outcomes of our sensors (Fig. 5C-E). This result could be attributed to potential variations from a combination of steps during the sampling and sensing processes. The observed trend differences might likely arise from inherent variability among sensors (though minimal variation was seen in calibration steps), the small sample size of cuboids, and potential variations in baseline viability. Nevertheless, despite these intrinsic variations, the sensor platform effectively captured the CytC production trends associated with different drug treatments. The result underlines the system’s potential as a promising platform for enhancing the accuracy of *ex vivo* anti-cancer drug screening on microdissected biopsies.

As we look toward the future, particularly in the context of long-term monitoring of CytC in culture using electrochemical aptasensors with slow *k*_off_ rates, several enhancements and considerations can further optimize the effectiveness and reliability of the sensor platform. It is essential to consider factors like sensor saturation points and the need for sensor regeneration, especially after calibration, for our slow dissociation rate. For instance, altering the aptamer sequence for lower binding affinity or modifying the sensor surface to amplify the signal response could be explored to refine the platform’s performance. This strategy holds promise not only for CytC but also for the development of aptamers targeting other relevant cancer biomarkers.^8–10^

Our sensing platform is an encouraging step in the real-time detection of CytC in long-term cultures of intact tumor biopsies. Key studies would further validate this approach. These include 1) the utilization of various biopsies from diverse patients; 2) the establishment of extended culture periods to enable cross-comparisons among multiple biomarkers, facilitating the accumulation of meaningful data; and 3) the comparison of our results with alternative approaches in patient studies. Nevertheless, the versatility of the presented technology allows for its adaptation to the multiplexing of a wide array of relevant biomarkers from human tissues and patient-derived samples, further advancing the real-time monitoring of cancer disease models and cancer drug screening platforms. The platform’s scalability allows for its future adaptation to multiplexed monitoring of diverse biomarkers, offers comprehensive insight into cellular responses, and enhances its applicability within intricate culture environments. These advancements will contribute to a more in-depth understanding of intact biopsy responses to combination therapies, thus expediting the development and implementation of enhanced treatment modalities for cancer.

## MATERIALS AND METHODS

### Regents and materials

All reagents were purchased from Thermo Fisher Scientific (MA, USA) unless stated otherwise. The recombinant human cytochrome c was purchased from RayBiotech Life, Inc (GA, USA). Cytochrome C aptamer (100 mM) and IDTE pH 7.5 (1X TE solution) were purchased from Integrated DNA Technologies Inc. (IA, USA). Methylene blue succinimidyl ester (MB-NHS) was purchased from Biotium (CA, USA). 6-Mercapto-1-hexanol (MCH) was purchased from TCI America Inc. (OR, USA). UltraPure DNase/RNase-free distilled water was purchased from Thermo Fisher Scientific (MA, USA). Phosphate-buffered saline powder (PBS; 10X, pH 7.4), Urea powder (Bioreagent), Sodium Chloride (ACS reagent), Nafion perifluorinated resin solution (5 wt. %), Gold (III) chloride hydrate (HAuCl4.xH2O, ∼50% Au basis) were purchase from Sigma Aldrich (MO, USA). PMMA was purchased from Astra Products, Inc. (NY, USA). Dry film photoresist (DF-1050) was generously given by Nagase ChemteX (OH, USA). Ag/AgCl (3M KCl) was purchased from BASi (IN, USA). Deionized water (DI) was purified using Milli-Q (Millipore, Bedford, MA). Drugs were diluted from DMSO stocks (10–200 mM), purchased from MedChem Express, and cisplatin (MedChemExpress, 3 M stock in DI water).

### Fabrication of microfluidic chip and integration of sensors on the platform

We utilized AutoCAD 2021 (Autodesk, CA, USA) to design the sensor pattern and the patterns for the sealing layer, connectors, insulation layer, loading frame, and a bottomless 24-well plate. Employing CO_2_ laser micromachining on PMMA, we fabricated all layers with different optimal laser powers and speeds. The CO_2_ laser system used (VLS3.60, Scottsdale, USA) has a wavelength of 10.6 μm and a maximum power of 30 W. Fig. S1 depicts the schematic for the sensor fabrication process. To fabricate the electrode (Fig. 1C), we generated a high-resolution photomask (10,160 DPI resolution, Fine Line Imaging, CO, USA) of the sensor pattern. Subsequently, we relied on dry film photoresist (Nagase ChemteX, OH, USA) as a sacrificial layer for Au deposition. We initiated this process by hot rolling the negative dry film photoresist (50 μm) onto the PMMA, employing a double-side hot roll laminator (SKY-335R6, Sky-Dsb Co., Seoul, KR). Afterward, we affixed the electrode pattern onto the dry film resist layer using a contact mask aligner (2001, Myriad Semiconductor, AR, USA). We exposed it to UV light with exposure energy ranging from 250-300 mJ/cm² for 30 s.

Following this, we engaged in a four-step development procedure: 1) Applying fresh running cyclohexanone through three cycles of 60 seconds each, 2) subjecting the samples to sonication in cyclohexanone for 1 min, 3) washing them with isopropanol for at least 1 mi, 4) rinsing them under running DI water. Subsequently, we dried the samples using an air gun and stored them in an airtight container before depositing Au using electron beam evaporation. To retain the option of removing the sacrificial layer post-Au deposition, we chose not to subject the photoresist layer to the final hard bake step. The next step involved treating the samples with oxygen plasma utilizing a reactive ion etching system (100 W, 75 mT, 80 sccm O_2_, 1 min). Subsequently, a Cr/Au (10/100 nm) layer was deposited through electron beam evaporation, after which the gold planar electrode array was formed by physically lifting the dry film resist sacrificial layer.

We manually applied 3M™ High-Strength Acrylic Adhesive 300LSE (MN, USA) to line the insulation layer (PMMA 0.5 mm thick), loading frame, and well layer (PMMA 6.4 mm thick, 1227T569, McMaster-Carr, Elmhurst, IL) before laser cutting (Fig. 1B). Next, we assembled the sensor layers along with the insulation layer and loading frame by removing the 3M300LSE liner and placing them under a heat press (Model 3912, Carver, IN, USA) at 200 psi for 5 minutes at room temperature. After this assembly, the sensors were subjected to cleaning and Ag/AgCl deposition steps. We followed the same procedure as mentioned above to assemble the well layer. However, this assembly step was only performed before the sensor was ready to be incubated with the CytC aptamer.

### Cleaning process for WE and RE surfaces

We performed a gold cleaning protocol employing a combination of H_2_O_2_ + KOH solution and KOH sweeping to clean the Au surface. For this purpose, we applied a solution of 50 mM KOH and 25% H_2_O_2_ to bare Au surfaces (including all surfaces intended for WE, RE, and CE) for 5 min (extending the duration could potentially lead to lifting off of the Au surface). Subsequently, we thoroughly rinsed the surfaces with DI water. Following the initial treatment, we subjected each electrode to a potential sweep ranging from 0.2 to 1.2 mV (vs. Ag/AgCl) at a scan rate of 50 mV/s in a solution of 50 nM KOH. Afterward, we rinsed the electrodes with DI water. A standard glass-bodied reference electrode with a porous junction (Ag/AgCl with 3M KCl) and a Pt wire were employed as the reference and counter electrodes, respectively. We also documented the AuNPs electrodeposition process in the supplementary method.

### Fabrication and characterization of Ag/AgCl film for the reference electrode

Once the Au surface had been cleaned, silver was electrodeposited onto the surface at -0.3 V for 300 seconds using a plating solution containing 250 mM silver nitrate, 750 mM sodium thiosulfate, and 500 mM sodium bisulfite.^76^ Subsequently, the electrode underwent treatment with 10% hydrochloric acid (HCl) for 1 minute. An aqueous solution comprising 0.1 M potassium chloride (KCl) and 0.01 M HCl was employed for anodic chlorination. Linear sweep voltammetry (LSV) was conducted, spanning the open circuit potential (OCP) to 0.4 V at a scan rate of 20 mV/s. This step was followed by ten cyclic voltammetry (CV) cycles within the 0.1 to 0.3 V range versus Ag/AgCl, employing a fixed scan rate of 100 mV/s. Subsequently, the electrode was thoroughly rinsed with deionized (DI) water and then incubated with 10 µL of 3.5 M KCl for 1 minute. The KCl droplet was subsequently removed, and the electrode was once again rinsed extensively with DI water. After drying the electrode with nitrogen (N_2_) gas, it was coated with a 10 µL droplet of 0.5% Nafion solution and left to dry overnight.^77^ A standard glass-bodied reference electrode with a porous junction (Ag/AgCl with 3M KCl) and a Pt wire were utilized as the reference and counter electrode, respectively (Fig. S5B).

### Modify sensor platform to become injection analysis platform for characterization of binding kinetics

We modified the existing platform by introducing an additional PMMA layer to establish channels and altering the well layer to create a sealed chamber for flow injection. The PMMA layer (0.3 mm thick) was lined with 3M300LSE adhesive before being precisely cut using a CO_2_ laser. We injected the culture medium through tubing (Tygon®, Cole-Parmer, IL, USA) connected to the injector loop at a 2.5 mL/min flow rate using a Fusion 200 syringe pump (Chemyx, Inc, TX, USA). Following a 5-minute equilibration period during which the sensors were exposed to continuous medium flow, we introduced CytC into the flow as the target compound.

### Cyt C aptamer sequences

The Cyt C aptamer sequence is 5ʹ-/ThioMC6-D/CC GTG TCT GGG GCC GAC CGG CGC ATT GGG TAC GTT GTT GC/AmMO/-3ʹ.^53,54^ Specifically, the DNA oligo was modified with a disulfide (S─S) bond at the 5ʹ terminus through a six-carbon (C6) spacer and an amino modifier at 3ʹ terminus. It was HPLC purified and arrived as lab ready (100 mM in IDTE, pH 8.0) and stored at -20 °C until used.

The redox reporters MB-NHS were conjugated to the 3’-terminus of amino-modified CytC aptamer through succinimide ester coupling following previously reported protocols.^9^ In short, MB-NHS (5 mg) was first dissolved in DMSO to a final concentration of 10 mg/ml and stored as aliquots in the dark at -20 °C. We added MB-NHS (10 mg/mL) with 100 mM aptamer solution so that NHS dye solution was 50 molar excess dye to aptamer ratio.^9^ The mixture was vortexed for 10 s to help disperse and allowed to react for 4 hrs at room temperature in the dark.

### Functionalization of aptamer on the working electrode

The working electrode was rinsed with copious deionized water (DI) and dried under N_2_ gas before aptamer immobilization. TCPE stock (100 mM) was prepared with UltraPure DNase/RNase-free distilled water that remained fresh for 3 months when stored in the dark at -20 °C. Conjugated MB-CytC aptamers were reduced by 100 mM TCPE with final concentration 1,000-fold of aptamer concentration at room temperature for 1 hour to cleave the S-S bond. Consecutively, the oligos were then diluted to 5 mM using TE buffer and vortexed for 10 s to help disperse. Five microliters of CytC aptamer (5 mM) solution were dropped on the working electrode and incubated in an airtight humidity petri dish at 4 °C overnight (∼15 hrs). The aptamer-immobilized electrodes were subsequently rinsed with copious DI water. MCH was diluted to 10 mM with UltraPure DNase/RNase-free distilled water before dropping onto each working electrode (5 mL) and incubated airtight at room temperature for at least 7 hrs. Lastly, the sensors were rinsed with copious DI water and stored in filter PBS (1X, pH 7.4) until used.

### Electrochemical signal measurement

All the electrochemical measurements were performed using a Digi-Ivy potentiostat (DY-2219, Digi-Ivy, Austin, TX, USA). For the SWV measurements, the sensors were integrated from 0.0 to -0.5 V (versus Ag/AgCl) with 40-mV amplitude and a voltage increment of 1 mV suitable frequency setting (ranging from 10 Hz to 1,000 Hz). A conventional three-electrode cell was used in electrodeposition, Ag/AgCl with 3M KCl as reference electrode, and platinum wire (0.05 mm diameter, Alfa Aesar, MA, USA) as counter electrode.

### Quantitative analysis and characterization of the Cyt C aptamer sensor

The CytC sensors were characterized in cultured medium DMEM-F12-10% FBS with spiked human CytC recombinant (reconstituted in PBS). The SWV measurement for aptasensors was scanned from -0.5 V to 0 V with 40-mV amplitude and a step potential of 1 mV. The frequency is 50 Hz and 1,000 Hz.

### Response of sensor for regeneration

Due to the slow *k*_off_ rate, our sensor platform requires regeneration after calibration before *in vitr*o and *ex vivo* testing. We explored various regeneration methods to identify one suitable for our platform.^78^ We applied multiple chemical regenerations with different pH, such as sodium hydroxide solution (0.05 M, pH 10), glycine in sodium hydroxide buffer (0.1 M, pH 8), and glycine in hydrochloric acid (0.1 M, pH 3.0). Additionally, a high-concentration sodium chloride solution (2M) was used to adjust ionic strength. High-concentration urea (6M, pH 7.0) was employed for regeneration, and thermal regeneration using PBS at 60 °C was also considered. As a result, although the signal gain differed slightly from the original due to the removal of some loosely bound aptamers, the regeneration with urea (6M) yielded the most favorable outcomes without compromising sensor functionality (Fig. S8).

### Hardware design

The primary elements consist of the analog switches and multiplexer, the RFduino ESP32 microcontroller, and the PCB circuitry required to interface with the sensor platform and multiplexing chips. We used Eagle 2022 (Autodesk, USA) to design the circuit schematics (Fig. S10). Subsequently, we created the component layout and wire connections for the two-layer PCB prototype, with one dedicated to interfacing with the sensor platform (Fig. S10A) and another for housing the multiplexing chips (Fig. S10B). Both designs were fabricated and partially assembled using a PCB manufacturing service (JLCPCB, HK). We connected the sensor platform to the PCB using standard pin connectors and established a connection between the PCB and the multiplexing housing board using FFC.

### Construction of multiplexing system

For our implementation, we utilized the RFduino ESP32 microcontroller (HiLetgo), which features a low-cost 32-bit LX6 microprocessor. The ESP32 chip package includes GPIO ports and an I2C bus for interfacing with peripheral components. Moreover, the ESP32 is compatible with the Arduino development environment. In order to achieve multichannel operations, we employed multiplexer chips. A multiplexer is a device that features multiple inputs, a single output, and control signals to determine which input is routed to the output. For our specific application, we utilized three analog-switched 32-to-1 multiplexer components (Analog Devices Inc., ADG732). These components have on-chip latches that facilitate the necessary hardware logic, allowing for control through GPIO. All WE, RE, and CE of the sensor array were connected to the corresponding WE, RE, or CE multiplexer chip. The multiplexer chips were connected in parallel to the ESP32’s +5V and GND pins for their power supply. GPIO pins on the ESP32 were connected to the 5 control pins on the multiplexer chips. These pins were configured to output either a +5V or 0V signal, corresponding to a 1 or a 0. The multiplexers then outputted a signal that depended on the configuration of the 5 input signals, as shown in the table displayed in Fig. S11 (ADG732 Datasheet, Analog Devices). The setup allows for quick and effortless shifting between different sensors on the array.

### Cell culture and drug screening

We grew U-87 MG cell (ATTC) in DMEM-F12 supplemented with 10% FBS and penicillin/streptomycin (Invitrogen). We conducted cell passages biweekly.

### Mouse tumor models and microdissection cuboid procedure

Mice were managed following institutional guidelines and under protocols endorsed by the Animal Care and Use Committee at the University of Washington (Seattle, USA). To create xenograft tumor models, male athymic nude mice (Jackson Laboratories, Foxn1nu) aged 6–10 weeks received subcutaneous injections in the flank region (1-2 million cells in 200 μL of serum and antibiotic-free DMEM-F12 medium). The mice were euthanized before the tumor volume exceeded 2 cm² (within 2–4 weeks).

The cuboidal-shaped microdissected tissues were processed using a previous protocol.^40^ In brief, we first cut the tissue into 400 μm-thick slices using a 5100 mz vibratome (Lafayette, Instrument) in an ice-cold HBSS solution (1X) and transferred the slices into ice-cold DMEM-F12 with 15 mM HEPES (Invitrogen). We cut the slices into cuboids using a standard Mcllwain tissue chopper (Ted Pella, Inc., USA) with two sets of orthogonal cuts separated by rotation of the sample holder. After separating the cuboids in DMEM-F12-HEPES by gently pipetting them up and down, we filtered them through a 750 μm and 300 μm filters (Pluriselect, USA). The cuboids were washed three times with DPBS, followed by a final fresh wash with DMEM-F12 containing HEPES. Subsequently, we cultured the cuboids in DMEM-F12-10% FBS overnight, allowing them to recover and remove any cell damage from handling before seeding them onto the sensors.

### Human colorectal cancer tumor supernatant

Colorectal tumor biopsy was obtained from a 43-year-old male with metastatic cancer who had been off treatment for a year.^40^ The cuboidal-shaped μDT tissues were processed using a previously established protocol as stated above.^40^ Subsequently, we cultured the cuboids under different drug conditions: FOLFOX (1 μg/mL 5FU + 1 μg/mL Oxaliplatin), FOLFIRI (1 μg/mL 5FU + 2 μg/mL Irinotecan), Staurosporine (1 μM), and a vehicle control (0.2% DMSO) for three days in modified William’s E Media.^79^

### Electrochemical recording from the supernatant of microdissected tumor tissue culture

We calibrated the sensor platform using human CytC recombinant ranging from 0 – 10,000 ng/mL. Following the calibration, the sensors are regenerated with 6 M urea for 5 min and stored in PBS at 4 °C for next-day measurement. We first conducted SWV measurements to collect the baseline signal, utilizing a potential range of -0.5 V to 0 V, with an amplitude of 4 mV and pulse frequencies set at 50 and 1,000 Hz. We added an appropriate volume of supernatant samples to achieve final concentrations that were 10-fold diluted compared to the stock solution.

### Encapsulation of microdissected tumor tissues using Cellink bioink

After the overnight recovery, we transferred the cuboids into a 96-well plate with 4 cuboids in each well to prepare for their seeding onto sensors. We transferred cuboids using a pipette with wide bore tips made by cutting off the ends. Cellink bioink, diluted to a 50% concentration with DPBS, was mixed well in a 1 mL Eppendorf tube. After removing the medium from the 96-well plate, we added 10 μL of the diluted Cellink bioink to each well and immediately transferred the encapsulated cuboids in bioink onto the WE surface. To crosslink the gel around the encapsulated tissues and ensure its mechanical stability on the WE, we introduced 150 μL of a 50 mM CaCl_2_ crosslinking agent (CELLINK, CA, USA) for 10 min. After three rounds of washing with DPBS, we added a growth medium, DMEM-F12-10% FBS, for subsequent culture.

### *Ex vivo* electrochemical recording from microdissected tumor tissues cultured with drugs

We calibrated the sensor platform using human CytC recombinant ranging from 0 – 25000 ng/mL in DMEM-F12-10% one day before the incubation. After calibration, the sensors are regenerated with 6 M urea for 5 min and stored in PBS at 4 °C for next-day incubation. We placed the encapsulated cuboid sensor platform in a 37 °C incubator for a minimum of 4 hrs to establish the baseline stability. Subsequently, we conducted SWV measurements to collect the baseline signal, utilizing a potential range of -0.5 V to 0 V, with an amplitude of 4 mV and pulse frequencies set at 50 and 1,000 Hz. Drugs were diluted in the cell culture medium for the drug treatment experiment and then transferred to the designated wells. We cultured the sensor platform in the 37 °C incubator during the experiment and removed it for measurement every 12 hrs over 48 hrs.

### Tissue processing and staining, brightfield microscopy, and image analysis

We stained live cuboids for 1 hr at 37 °C using SYTOX green (SG; Invitrogen, 0.01 μM) with or without Hoechst (H; Invitrogen, 16 μM). Epifluorescence and brightfield microscopy of the cuboids were performed using a BZ-X800 microscope (Keyence, IL, USA) at 2× magnification. We utilized FIJI software to analyze SG intensity as follows. We first subtracted the background fluorescence determined from empty areas. We then established cuboid regions from the Hoechst channels using thresholding and watershed techniques on a binary image. With the Analyze Particles function, we acquired the mean SG fluorescence, which we subsequently normalized to the average value of untreated cuboids.

### Electrochemical data analysis

The frequency of the applied square wave influences the signaling of the aptasensor. Square wave voltammetry can be adjusted to result in either an increase (Signal-ON) or a decrease (Signal-OFF) in peak current upon the introduction of the target molecule (Fig. 2E). To address signal drift and improve signal strength during *ex vivo* measurements, voltammograms are captured at two different frequencies, and these are then transformed into "Kinetic Differential Measurement" (KDM) values.^4,59^ These KDM values are calculated by subtracting the normalized peak currents observed at the Signal-ON and Signal-OFF frequencies and dividing them by their average. To create a calibration curve, the averaged KDM values obtained *in vitro* across a range of target concentrations are fitted to a Hill-Langmuir isotherm equation (Eq. 1)^4,59^. The calibration curve fitting enables the extraction of all the parameters nH, K_1/2_, KDM_min_, and KDM_max_.

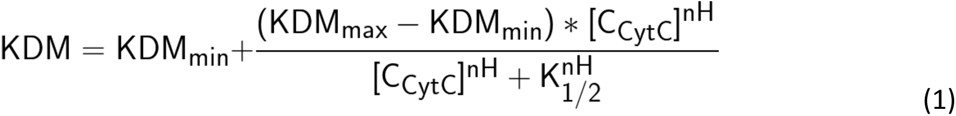

nH represents the Hill coefficient, which measures the degree of binding cooperativity.

K_1/2_ denotes the midpoint of the binding curve.

KDM refers to the observed KDM value at the given target concentration.

KDM_min_ represents the KDM value recorded in the absence of the target.

KDM_max_ represents the anticipated KDM value when the target is fully saturated.

C_CytC_ refers to the concentration of interested analyte (CytC).

Once derived from a calibration curve, these parameters facilitate the conversion of aptasensor output into estimations of the target concentration (Eq. 2):^4,59^

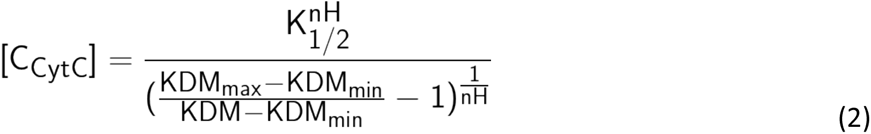

### Statistical analyses

We used GraphPad Prism 10 (Boston, MA, USA) to perform statistical tests and data visualization. We used MATLAB (R2021a, USA) to analyze all experiment data. First, we exported the raw data to MATLAB, extracted the MB redox peak currents from record voltammograms and calculated KDM values. Subsequently, we extracted the MB redox peak currents from recorded voltammograms to compute KDM values. We entered titration concentration and average KDM values into Eq. 1 in MATLAB to fit the data to a Hill-Langmuir isotherm. We fitted the data to Eq. 1 with the following constraints: -∞ < KDM_max_ < ∞, 1×e^−1^^2^ < K_1/2_ < 1, -∞ < KDMmin < ∞, 0 < nH < 10.^4,59^

## Supplementary Materials

Fig. S1. Dry film resist fabrication process of microelectrodes.

Fig. S2. Schematic and cross-section of modified sensor platform with loop injection system.

Fig. S3. KDM response of CytC in phosphate-buffer saline.

Fig. S4. The characterization of Au-aptasensor vs. AuNPs-aptasensors and their KDM signals.

Fig. S5. Characterization of the Au planar surface cleaning process and the Ag/AgCl surface fabrication for the reference electrode.

Fig. S6. Characterization of aptasensors response through multiplexers.

Fig. S7. Characterization of the response signal of aptasensors with Cellink bioink and collagen at 50 Hz and 1,000 Hz.

Fig. S8. Response signal of aptasensors after regeneration with 6 M urea.

Fig. S9. Selectivity of the CytC aptasensors to relevant drugs.

Fig. S10. Circuit diagram of multiplexing housing chips PCB and sensor interfacing PCB.

Fig. S11. Specification of the multiplexer chip ADG732 datasheet.

Data S1. (separate file). A downloadable file for the circuit diagram of the multiplexing chips housing PCB. Data S2. (separate file). A downloadable file for the circuit diagram of the sensor interfacing PCB.

## Acknowledgments

We thank Kim A. Woodrow’s group for the use of their microplate reader.

## Author Contributions

T.N.H.N., L.H., T.K., R.S., N.A.C., A.F. designed experiments. T.N.H.N., L.H., T.K., E.L., H.K. performed experiments. T.N.H.N. performed analysis. T.N.H.N., L.H., N.A.C., A.F. wrote the manuscript.

## Data and Materials availability

All data needed to evaluate the conclusions in the paper are present in the paper and/or the Supplementary Materials.

