## Supplementary Files for "Label-Free, Real-Time Monitoring of Cytochrome C Responses to Drugs in Microdissected Tumor Biopsies with a Multi-Well Aptasensor Platform"

**This PDF file includes:**

Figs. S1 to S11
Data S1 to S2
Supplementary Text

**Other Supplementary Materials for this manuscript include the following:**
Data S1 to S2

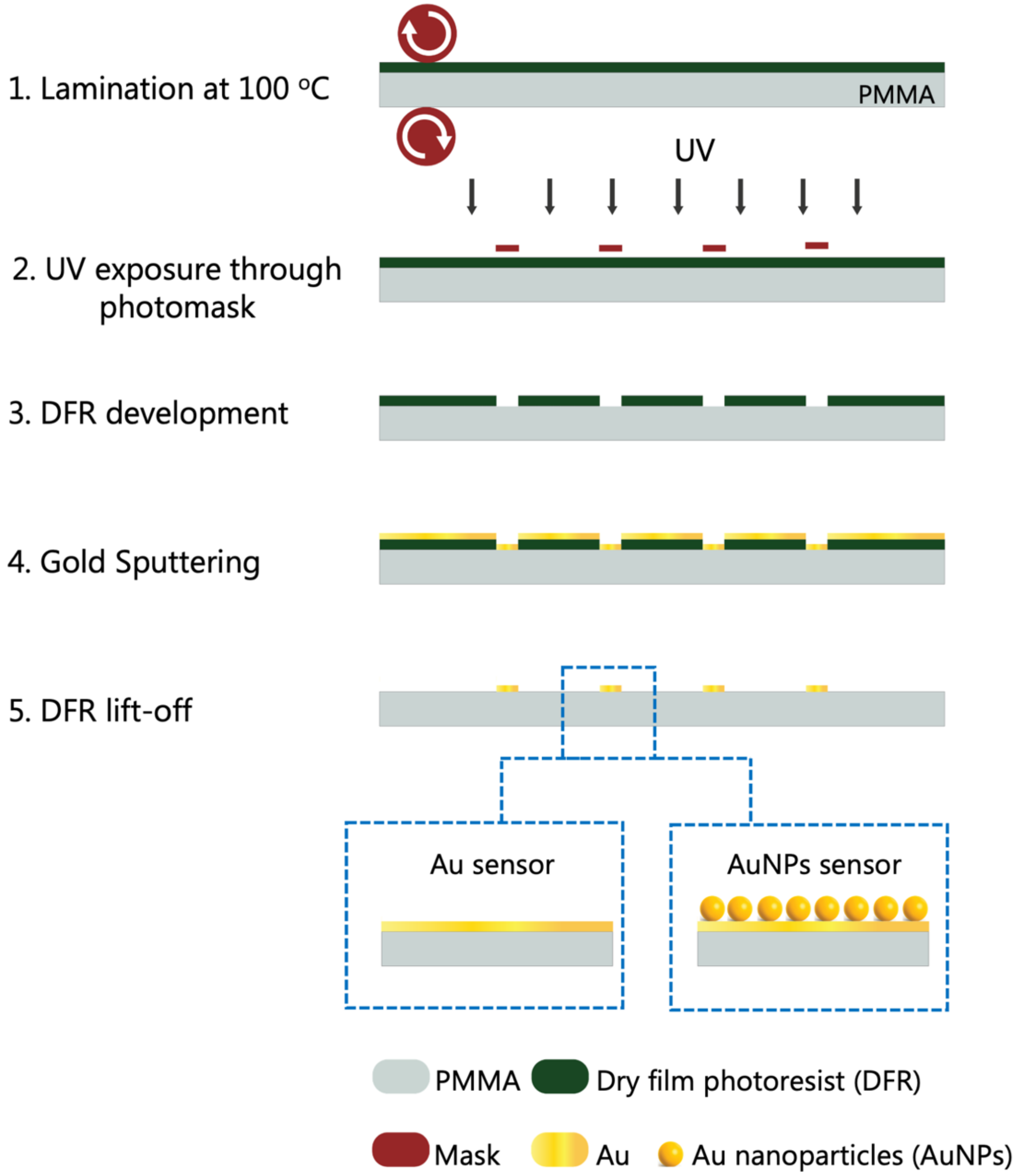


**Fig. S1. Dry film resist fabrication process of microelectrodes.** Dry film resist (DFR) is laminated and utilized as a sacrificial layer for Au deposition.

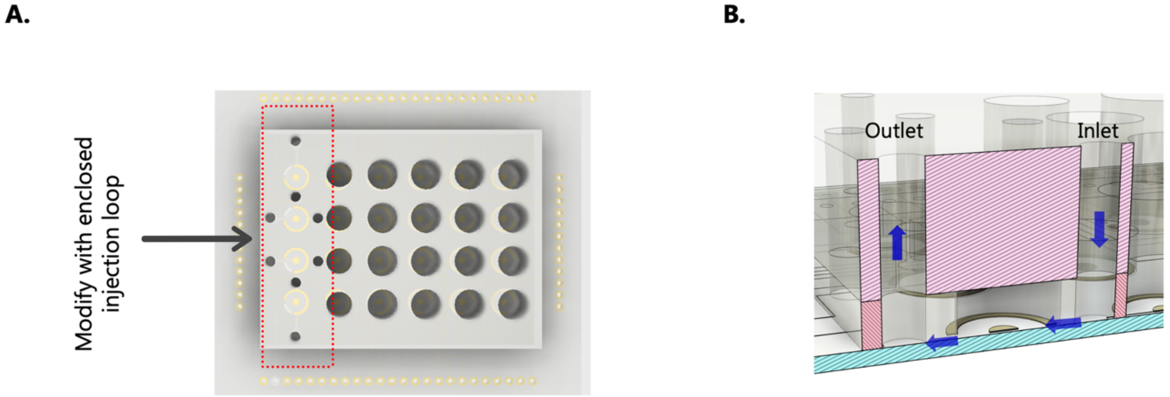


**Fig. S2. Modified sensor platform and selectivity testing. A.** Modified sensor platform with an inlet and outlet loop injection system**. B.** Cross-section of the platform. We modified the original platform (Fig. 1B) by incorporating an additional layer of $CO_2$ laser micromachined PMMA between the insulation and loading frame layers. This modification created small channels running across the sensor, forming an injection loop. Simultaneously, we adapted the well layer, closing the top to establish an enclosed injection loop.

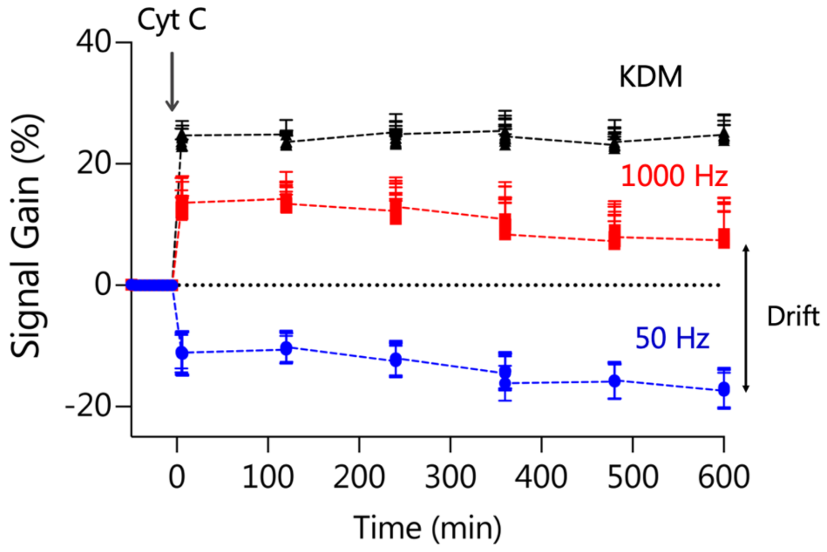


**Fig. S3. KDM response of CytC in Phosphate-buffered Saline (PBS).** KDM (black), as calculated from responses to CytC measured at 50 Hz and 1,000 Hz (blue and red lines, respectively), showed reduced drift over time and increased signal amplitude. Error bars represent the standard deviation (n = 3 sensors).

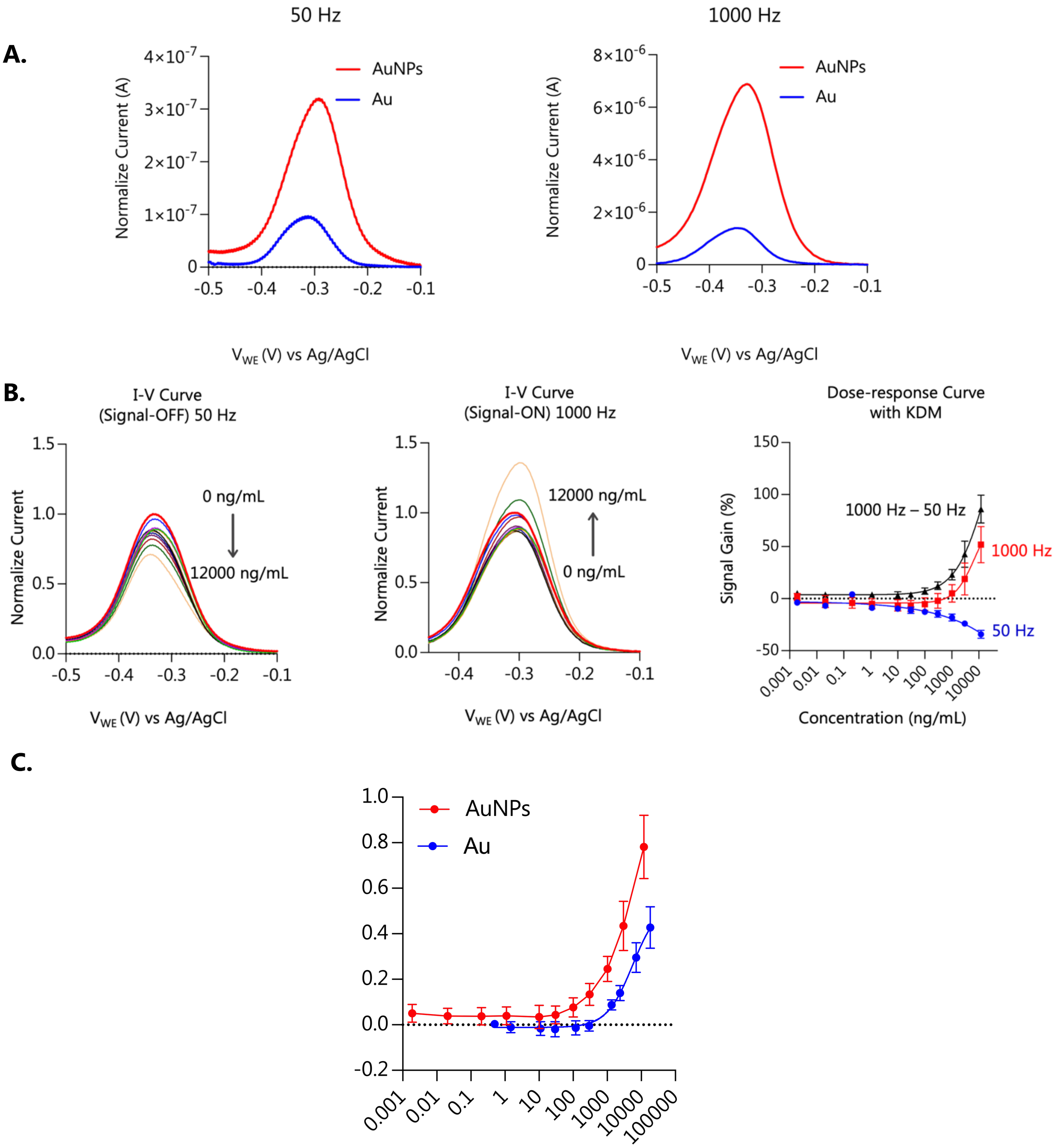


**Fig. S4. Signal comparisons between gold (Au) and gold nanoparticles (AuNPs) electrochemical aptasensors. A.** Square wave voltammogram of Au and AuNPs electrochemical aptasensors at 50 Hz and 1,000 Hz. **B. Responses of CytC aptasensors calibrated in DMEM-F12-10% FBS culture medium.** Response of AuNPs-aptasensors to CytC square wave voltammograms obtained for the 1 – 12,000 ng/mL concentration range at 50 Hz and 1,000 Hz (representative sensor, the arrow indicates an increase in concentration). Graph of KDM values calculated from the normalized peak height at 50 Hz and 1,000 Hz. Error bars represent the standard deviation (n = 5). **C.** Comparison of KDM responses between AuNPs-aptasensors and Au-aptasensors for concentrations that overlap from 0.1 ng/mL – to 12,000 ng/mL concentration range. Error bars represent the standard deviation (n = 5).

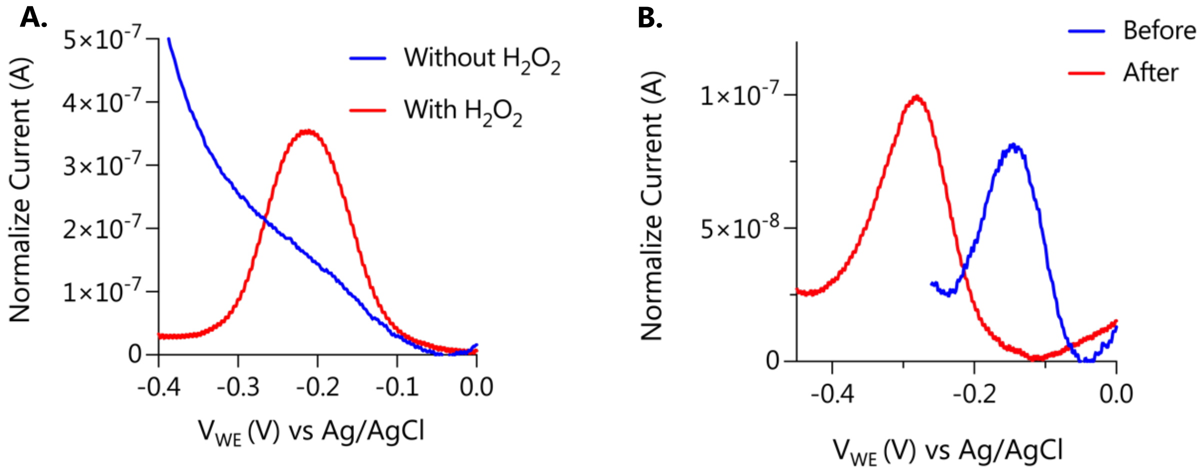


**Fig. S5. Electrochemical characterization of the Au-aptasensor system before and after modified cleaning and fabrication processes. A.** Square wave voltammogram of Au-aptasensors treated with $H_2O_2$ and without $H_2O_2$ before aptamer immobilization. **B.** Square wave voltammogram of Au-aptasensor system with the reference electrode before and after modified cleaning and additional fabrication processes.

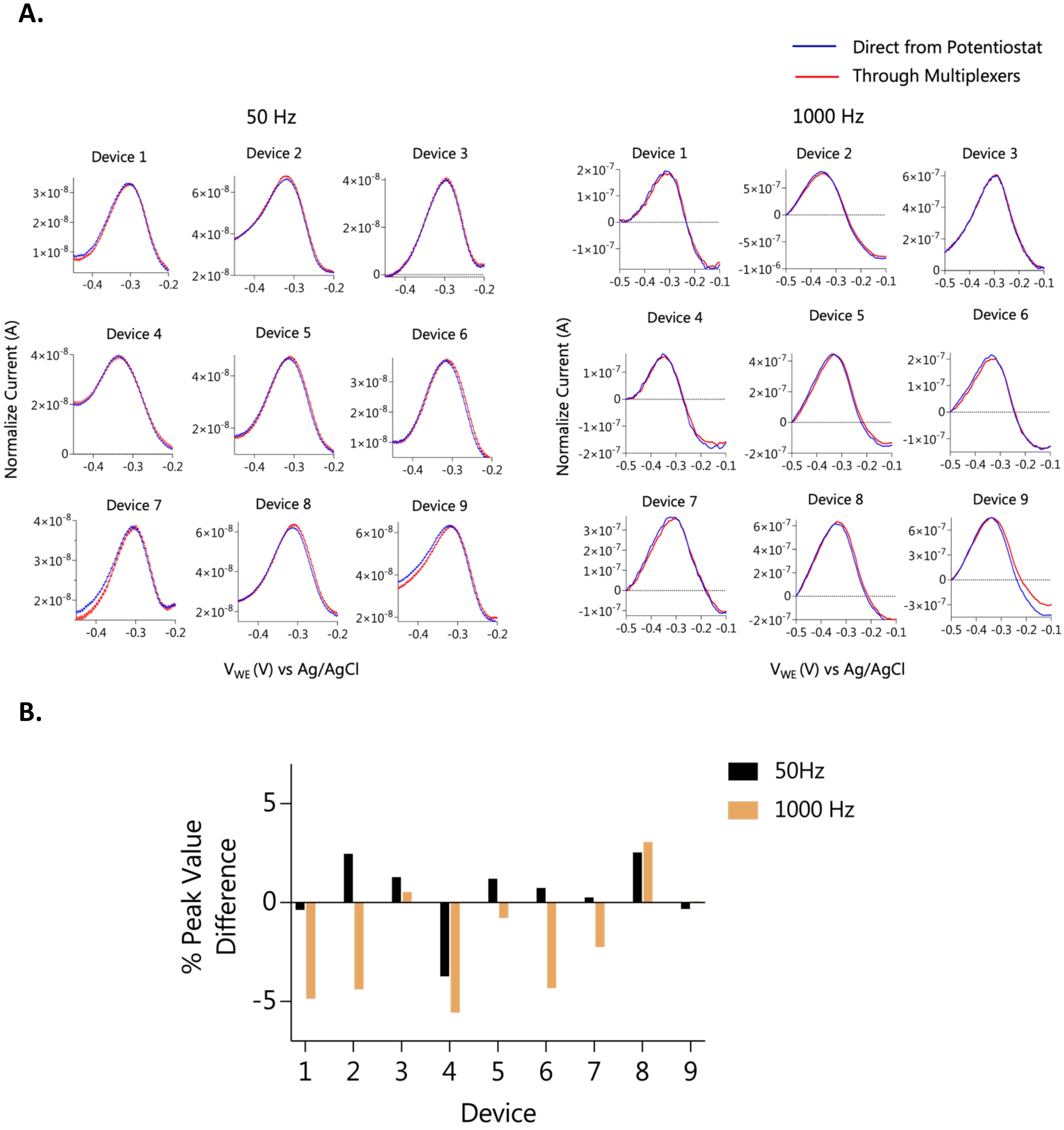


**Fig. S6. Signal response comparison with and without multiplexer. A.** The comparison of the square wave voltammogram of Au-aptasensors obtained from the sensor platform is connected directly through the potentiostat or indirectly through multiplexers. **B.** The comparison of % peak difference at 50 Hz and 1,000 Hz between sensor platforms connected directly to the potentiostat and indirectly to the multiplexer. The comparison % peak difference is less than 5% for both frequencies (50 Hz and 1,000 Hz).

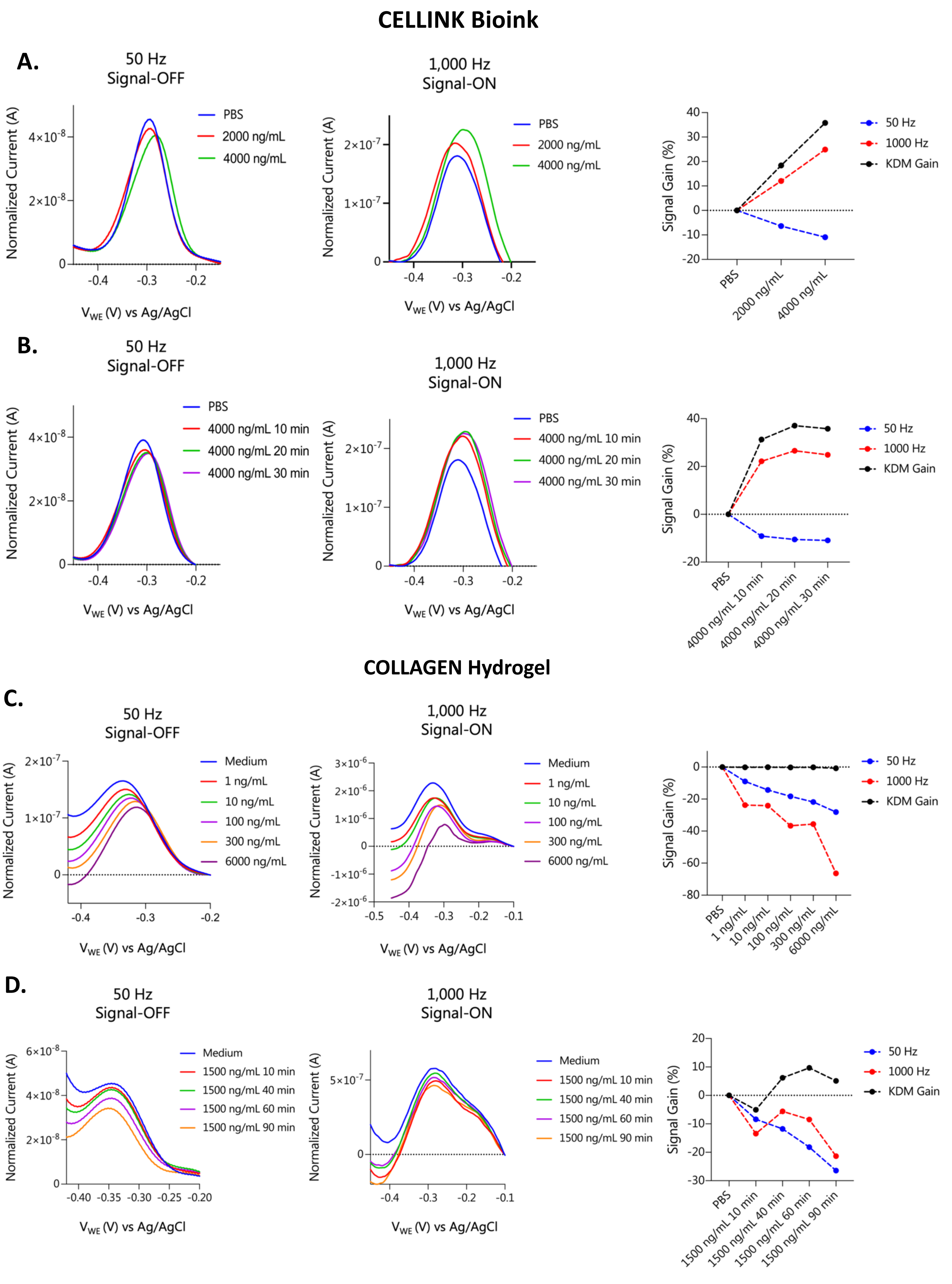

CELLINK Bioink
A.
50 Hz
Signal-OFF
PBS
2000 ng/mL
4000 ng/mL
Normalized Current (A)
V_WE (V) vs Ag/AgCl
1,000 Hz
Signal-ON
Signal Gain (%)
50 Hz
1000 Hz
KDM Gain
B.
50 Hz
Signal-OFF
PBS
4000 ng/mL 10 min
4000 ng/mL 20 min
4000 ng/mL 30 min
1,000 Hz
Signal-ON
COLLAGEN Hydrogel
C.
50 Hz
Signal-OFF
Medium
1 ng/mL
10 ng/mL
100 ng/mL
300 ng/mL
6000 ng/mL
1,000 Hz
Signal-ON
D.
50 Hz
Signal-OFF
Medium
1500 ng/mL 10 min
1500 ng/mL 40 min
1500 ng/mL 60 min
1500 ng/mL 90 min
1,000 Hz
Signal-ON

**Fig. S7. Characterization of sensor response with Cellink bioink and Collagen encapsulation. A.** Square wave voltammogram of Au-aptasensor responses to different CytC concentrations at 50 Hz and 1,000 Hz with **Cellink bioink** on the working electrode surface, and the signal gain responses corresponding to different CytC concentrations at 50 Hz, 1,000 Hz, and calculated KDM gain. **B.** Square wave voltammogram of Au-aptasensor responses to 4,000 ng/mL CytC at 50 Hz and 1,000 Hz with **Cellink bioink** over time, and the signal gain responses corresponding at 50 Hz, 1,000 Hz, and calculated KDM gain. **C.** Square wave voltammogram of Au-aptasensor responses to different CytC concentrations at 50 Hz and 1,000 Hz with **collagen hydrogel** on the working electrode surface, and the signal gain responses corresponding to different CytC concentrations at 50 Hz, 1,000 Hz, and calculated KDM gain. **D.** Square wave voltammogram of Au-aptasensor responses to 1,500 ng/mL CytC at 50 Hz and 1,000 Hz with **collagen hydrogel** over time, and the signal gain responses corresponding at 50 Hz, 1,000 Hz, and calculated KDM gain. Collagen encapsulation obstructed the aptasensor's ability to perform KDM and led to a prolonged reaction time, preventing the sensor from reaching an equilibrium state even after 90 minutes. We hypothesize that the density of the collagen gel partially impairs the conformational change of the aptasensor upon binding. Alternatively, Cellink bioink exhibited successful sensor performance.

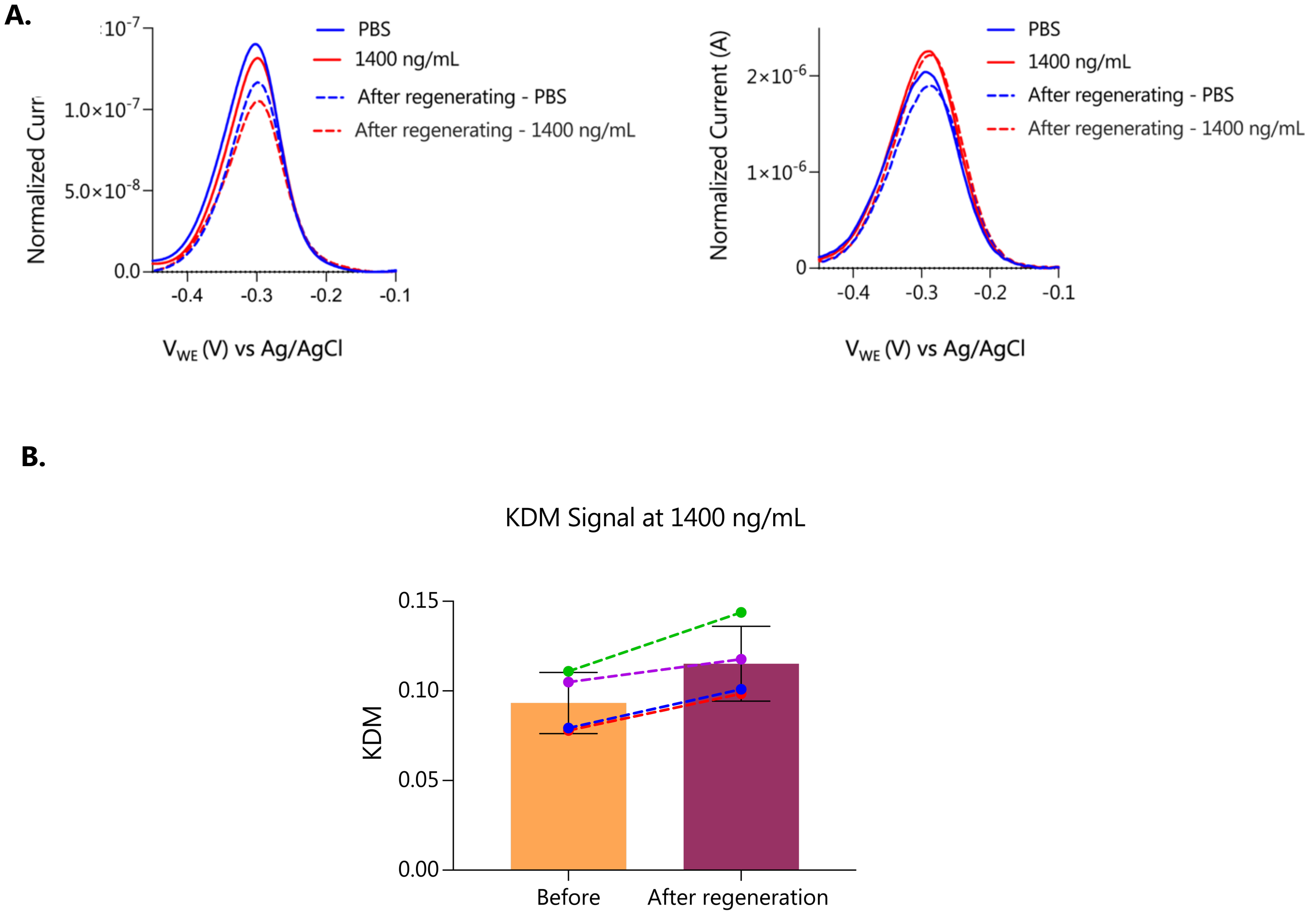


**Fig. S8. Signal recovery after regeneration. A.** Recovery of sensor signal after testing in culture medium at 50 Hz and 1,000 Hz. The dashed lines represent the recovery peaks obtained after washing with 6 M urea. **B.** The % of KDM signal change responses to 1400 ng/mL. The standard deviation from four independent sensors is presented here (n = 4).

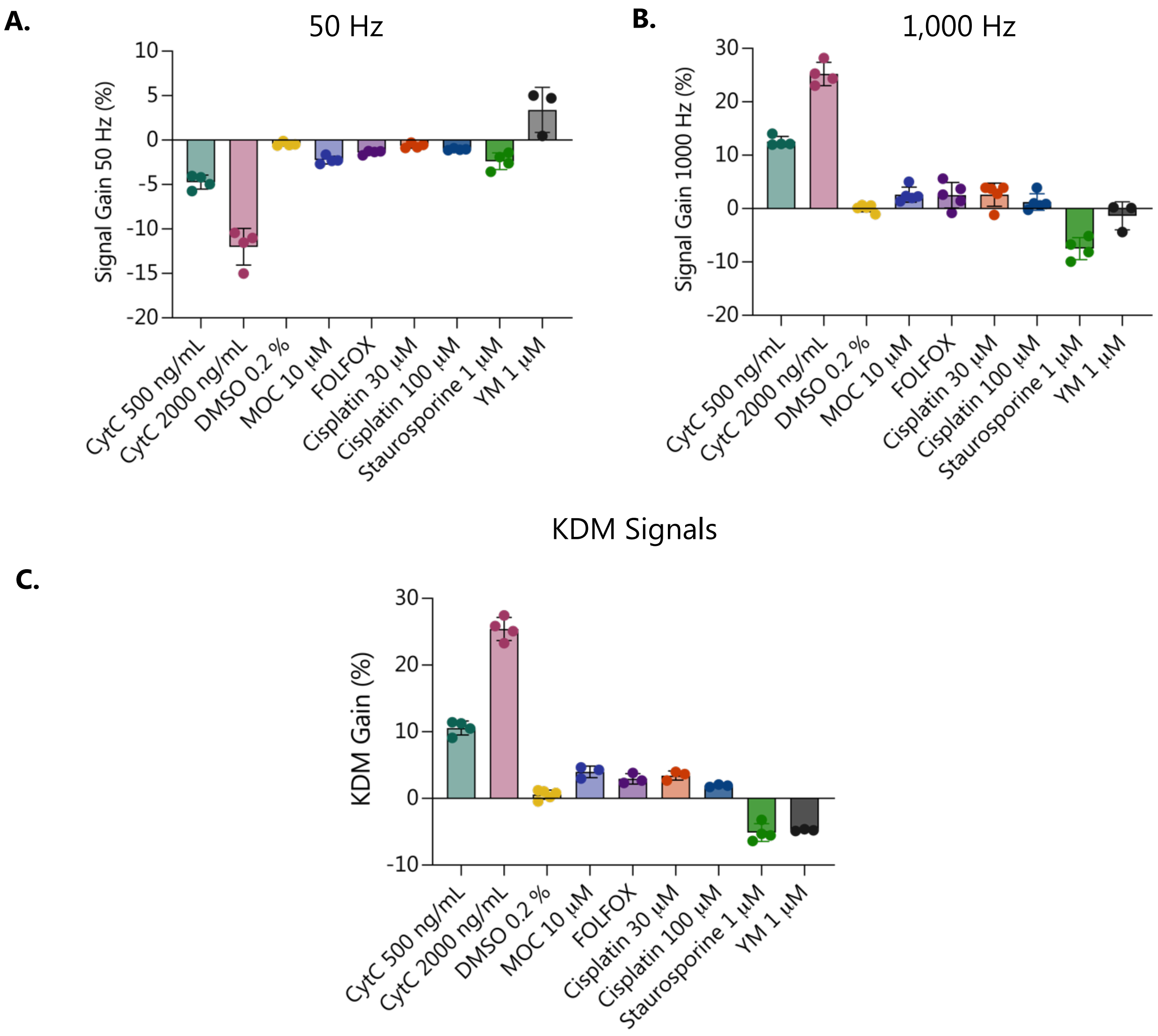


**Fig. S9. Selectivity of CytC aptasensors response to CytC vs. response to relevant drugs.** Response of CytC aptasensors to CytC (500 ng/mL = 8.3 nM and 2000 ng/mL = 166 nM) and to the drugs that we used to treat the microdissected tumor tissues, DMSO control vehicle (0.2 %), 10 µM MOC, FOLFOX (1 µg/mL 5FU + 1 µg/mL Oxaliplatin), 30 µM and 100 µM Cisplatin, 1 µM Staurosporine, and 1 µM YM-155. **A.** Measurements were performed in a culture medium at 50 Hz. **B.** Measurements were performed in a culture medium at 1,000 Hz. **C.** KDM signals were calculated from responses measured at 50 Hz and 1,000 Hz. Under these conditions, responses to CytC lead to a higher signal gain than responses to the relevant drugs.

A. B.

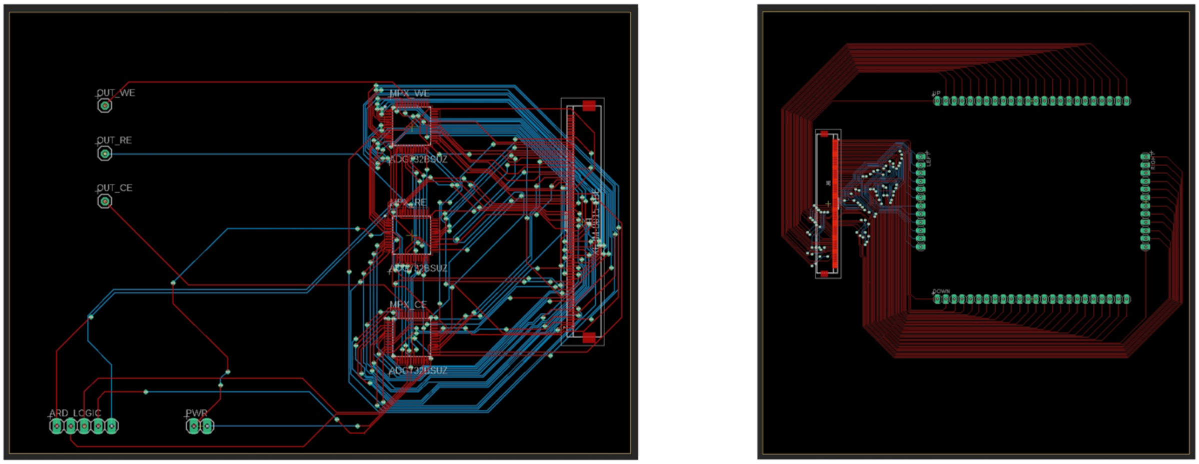


**Fig. S10. Multiplexer circuit diagram. A.** Circuit diagram for the multiplexing chips housing PCB. **B.** Circuit diagram for the sensor interfacing PCB.

**Data S1. (separate file)**

A downloadable file for the circuit diagram of the multiplexing chips housing PCB.

**Data S2. (separate file)**

A downloadable file for the circuit diagram of the sensor interfacing PCB.

### SPECIFICATIONS

#### +5 V SINGLE SUPPLY

$V_{DD}$ = 5 V ± 10%, $V_{SS}$ = 0 V, GND = 0 V, unless otherwise noted.

**Table 1.**

| Parameter | Symbol | ADG726/ADG732 | | ADG732 | Unit | Test Conditions/Comments |
|---|---|---|---|---|---|---|
| | | +25°C | −40°C to +85°C | −40°C to +125°C | | |
| ANALOG SWITCH | | | | | | |
| Analog Signal Range | | | 0 V to $V_{DD}$ | | V | |
| On Resistance | $R_{ON}$ | 4 | 5 | | Ω typ | $V_S$ = 0 V to $V_{DD}$, $I_{DS}$ = 10 mA, see Figure 20 |
| | | 5.5 | 6 | 7 | Ω max | |
| On Resistance Match Between Channels | $\Delta R_{ON}$ | | 0.3 | | Ω typ | $V_S$ = 0 V to $V_{DD}$, $I_{DS}$ = 10 mA |
| | | | 0.8 | 1 | Ω max | |
| On Resistance Flatness | $R_{FLAT (ON)}$ | 0.5 | | | Ω typ | $V_S$ = 0 V to $V_{DD}$, $I_{DS}$ = 10 mA |
| | | | 1 | 1.2 | Ω max | |
| LEAKAGE CURRENTS | | | | | | $V_{DD}$ = 5.5 V |
| Source Off Leakage | $I_S$ (Off) | ±0.01 | | | nA typ | $V_D$ = 4.5 V/1 V, $V_S$ = 1 V/4.5 V, see Figure 21 |
| | | ±0.25 | ±1 | ±2 | nA max | |
| Drain Off Leakage | $I_D$ (Off) | ±0.05 | | | nA typ | $V_D$ = 4.5 V/1 V, $V_S$ = 1 V/4.5 V, see Figure 24 |
| ADG726 | | ±0.5 | ±2.5 | | nA max | |
| ADG732 | | ±1 | ±5 | ±10 | nA max | |
| Channel On Leakage | $I_D$, $I_S$ (On) | ±0.05 | | | nA typ | $V_D$ = $V_S$ = 1 V, or 4.5 V, see Figure 25 |
| ADG726 | | ±0.5 | ±2.5 | | nA max | |
| ADG732 | | ±1 | ±5 | ±10 | nA max | |
| DIGITAL INPUTS | | | | | | |
| Input High Voltage | $V_{INH}$ | | 2.4 | 2.4 | V min | |
| Input Low Voltage | $V_{INL}$ | | 0.8 | 0.8 | V max | |
| Input Current | | | | | | |
| Low or High | $I_{INL}$ or $I_{INH}$ | 0.005 | | | μA typ | $V_{IN}$ = $V_{INL}$ or $V_{INH}$ |
| | | | ±0.5 | ±0.5 | μA max | |
| Digital Input Capacitance | $C_{IN}$ | 5 | | | pF typ | |
| DYNAMIC CHARACTERISTICS[1] | | | | | | |
| Transition Time | $t_{TRANSITION}$ | 23 | | | ns typ | $R_L$ = 300 Ω, $C_L$ = 35 pF, see Figure 27 |
| | | 34 | 40 | 48 | ns max | $V_{S1}$ = 3 V/0 V, $V_{S32}$ = 0 V/3 V |
| Break-Before-Make Time Delay | $t_D$ | 18 | | | ns typ | $R_L$ = 300 Ω, $C_L$ = 35 pF; $V_S$ = 3 V, see Figure 28 |
| | | | 1 | 1 | ns min | |
| On Time ($\overline{CS}$, $\overline{WR}$) | $t_{ON}$ ($\overline{CS}$, $\overline{WR}$) | 18 | | | ns typ | $R_L$ = 300 Ω, $C_L$ = 35 pF; $V_S$ = 3 V, see Figure 29 |
| | | 25 | 32 | 38.5 | ns max | |
| Off Time ($\overline{CS}$, $\overline{WR}$) | $t_{OFF}$ ($\overline{CS}$, $\overline{WR}$) | 17 | | | ns typ | $R_L$ = 300 Ω, $C_L$ = 35 pF; $V_S$ = 3 V, see Figure 29 |
| | | 23 | 29 | 33 | ns max | |
| On Time ($\overline{EN}$) | $t_{ON}$ ($\overline{EN}$) | 24 | | | ns typ | $R_L$ = 300 Ω, $C_L$ = 35 pF; $V_S$ = 3 V, see Figure 30 |
| | | 32 | 40 | 43 | ns max | |
| Off Time ($\overline{EN}$) | $t_{OFF}$ ($\overline{EN}$) | 16 | | | ns typ | $R_L$ = 300 Ω, $C_L$ = 35 pF; $V_S$ = 3 V, see Figure 30 |
| | | 22 | 25 | 25 | ns max | |
| Charge Injection | $Q_{INJ}$ | 5 | | | pC typ | $V_S$ = 2.5 V, $R_S$ = 0 Ω, $C_L$ = 1 nF, see Figure 31 |
| Off Isolation | $I_{SO}$ | −72 | | | dB typ | $R_L$ = 50 Ω, $C_L$ = 5 pF, f = 1 MHz, see Figure 22 |
| Channel-to-Channel Crosstalk | $C_{TK}$ | −72 | | | dB typ | $R_L$ = 50 Ω, $C_L$ = 5 pF, f = 1 MHz, see Figure 23 |
| −3 dB Bandwidth | BW | | | | | $R_L$ = 50 Ω, $C_L$ = 5 pF, see Figure 26 |
| ADG726 | | 34 | | | MHz typ | |
| ADG732 | | 18 | | | MHz typ | |

**Fig. S11. Specification of the multiplexer chips ADG732 datasheet.**

**Supplementary Text**

**Supplementary Method**

**Gold surface cleaning and fabrication of reference electrode silver/silver chloride surface.** In most cases, it is essential to have a high-quality electrode surface, such as bare Au, to provide the strong covalent binding of aptamers and efficiently collect the transduced signal for robust aptamer based sensing. Numerous studies have presented standard cleaning protocols utilizing repeated cyclic voltammetry (CV) with sulfuric acid ($H_2SO_4$).[1,2] We also recognized the significance of this cleaning step prior to AuNPs deposition or to our bare Au surface for aptamer immobilization.[3,4] However, due to the fabrication of our sensors on PMMA, conventional cleaning methods are not applicable. Published protocols typically employ solvents that do not affect glass but can damage plastics such as PMMA.[1,2]

As a result, we adopted an alternative cleaning protocol employing hydrogen peroxide ($H_2O_2$) and linear sweep voltammetry (LSV) in potassium hydroxide (KOH). This protocol, compatible with our platform, produces a high-quality Au surface suitable for subsequent processing.[5] We characterized the surface before and after cleaning. The cleaned sensor exhibited a distinct MB reduction peak with a flat baseline on AuNPs and bare Au surfaces. This observation indicated the formation of a compact self-assembled monolayer, in contrast to the surfaces of the uncleaned sensor. The MB reduction peak on these surfaces is hardly distinguishable from the background (Fig. S5A).

Our three-electrode system utilized an on-board reference electrode for each well. The generation of a stable Ag/AgCl film is essential to ensure mechanical stability for on-board real-time monitoring in the culture medium. An inherent limitation of thin film Ag/AgCl is its inadequate film adhesion and structural instability. Notably, a prevalent challenge encountered with Ag/AgCl film reference electrodes is the continual dissolution of $Cl^-$ ions, leading to detachment from the substrate.[6–8] This phenomenon impacts overall sensitivity and diminishes the electrode's lifespan. Considering our sensors' fabrication on PMMA, traditional methods for generating an AgCl surface (like prolonged exposure to high-concentration bleach solutions or chlorination process at high voltage/current)[9–11] are not viable.

Consequently, we adopted an alternative protocol to establish a thin yet robust Ag/AgCl/Nafion electrode surface that increases stability and rapid response.[6] This approach included Ag plating, chemical cleansing, cyclic chlorination, enhanced interfacial adhesion, and improved surface uniformity of the AgCl film. Moreover, introducing a final Nafion layer, a widely used cation-exchange polymeric membrane, extended the quasi-stable period of the Ag/AgCl while minimizing thickness (Fig. S5B).

**AuNPs electrodeposition.** To create an AuNP electrode, we employed a modified protocol based on a previous study.[12] The process involved immersing the Au electrode in a solution containing 2.5 mM $HAuCl_4$ and a 0.2 M $H_2SO_4$ aqueous solution. Before starting the experiment, an $H_2SO_4$

solution was purged of oxygen using high-purity nitrogen gas for 30 min to eliminate oxygen. Subsequently, we added $HAuCl_4$ into the deoxygenated $H_2SO_4$ solution and sonicated it in an ice bath until complete dissolution (~5 h). The electrodeposition of AuNPs was carried out for 300 seconds at -0.2 V (versus Ag/AgCl). We maintained the solution's temperature at 4 °C to achieve smaller particles using an ice bath. The resulting AuNP electrodes were thoroughly rinsed with DI water, dried using a stream of $N_2$ gas, and stored in airtight containers until ready for use.
